# Tissue-wide Effects Overrule Cell-intrinsic Gene Function in Cortical Projection Neuron Migration

**DOI:** 10.1101/2022.02.16.480659

**Authors:** Andi H. Hansen, Florian M. Pauler, Michael Riedl, Carmen Streicher, Anna Heger, Susanne Laukoter, Christoph Sommer, Armel Nicolas, Björn Hof, Li Huei Tsai, Thomas Rülicke, Simon Hippenmeyer

## Abstract

The mammalian neocortex is composed of diverse neuronal and glial cell classes that broadly arrange in six distinct laminae. Cortical layers emerge during development and defects in the developmental programs that orchestrate cortical lamination are associated with neurodevelopmental diseases. The developmental principle of cortical layer formation is based on concerted radial projection neuron migration, from their birthplace to their final target position. Radial migration occurs in defined sequential steps that are regulated by a large array of signaling pathways. However, based on genetic loss-of-function experiments, most studies have thus far focused on the role of cell-autonomous gene function. Yet, cortical neuron migration *in situ* is a complex process and migrating neurons traverse along diverse cellular compartments and environments. The role of tissue-wide properties and genetic state in radial neuron migration is however not well understood. Here, we utilized Mosaic Analysis with Double Markers (MADM) technology to either sparsely or globally delete gene function followed by quantitative single cell phenotyping. The MADM-based gene ablation paradigms in combination with computational modeling demonstrated that global tissue-wide effects predominate cell-autonomous gene function albeit in a gene-specific manner. Our results thus suggest that the genetic landscape in a tissue critically impacts the overall migration phenotype of individual cortical projection neurons. In a broader context our findings imply that global tissue-wide effects represent an essential component of the underlying etiology associated with focal malformations of cortical development (FMCD) in particular, and neurological diseases in general.

**LAY SUMMARY:** The assembly of the mammalian brain is a complex process and depends on tightly regulated developmental and genetic programs. The coordinated process of nerve cell (neuron) migration is essential to home neurons into their correct position which is vital for normal brain development. Disturbance of the neuron migration process at any point of development leads to severe brain malformations which result in devastating disease. To date, studies have mainly focused on cell-intrinsic gene functions controlling neuronal migration. Therefore, very little is known about the possible contribution of the cellular surrounding and tissue-wide effects. The scale and nature of such global tissue-wide effects remain completely unclear. We thus established genetic platforms based on Mosaic Analysis with Double Markers (MADM) to visualize and quantify tissue-wide effects in a defined genetic context and at single cell resolution. We found a critical predominant role of the genetic landscape and state of cellular environment affecting overall neuron migration properties. In a broader context our results suggest that global tissue-wide effects are a major component of the underlying etiology of neurological diseases such as focal malformations of cortical development (FMCD).

## INTRODUCTION

The mammalian neocortex is arranged in a six-layered structure and executes essential higher order cognitive brain functions. The laminated organization emerges during embryonic development and instructs the wiring diagram of cortical microcircuits (Hanganu-Opatz et al., 2021; Lodato and Arlotta, 2015). Deficits in the developmental programs orchestrating cortical layering represent a key underlying mechanism of neurodevelopmental disorders including cortical malformation (Guerrini and Parrini, 2010; Heng et al., 2010; Juric-Sekhar and Hevner, 2019).

A cardinal feature of cortical layering during development is the temporally sequential arrangement of the distinct layers in an ‘inside-out’ fashion (Angevine and Sidman, 1961; McConnell, 1995; Valiente and Marín, 2010). As such, progressively later born, radial glial cell (RGC)-derived, projection neurons migrate out radially past earlier born cohorts to gradually build up the inversely arranged laminae. Seminal live-imaging experiments at single cell resolution revealed that radial migration of cortical projection neurons occurs in a defined sequential sequence (Nadarajah et al., 2001; Noctor et al., 2001, 2004; Tsai and Vallee, 2011). First, newly-born projection neurons acquire a bipolar morphology and migrate from the ventricular zone (VZ) to the subventricular zone (SVZ) where they adopt a multipolar shape (Tabata and Nakajima, 2003). While in the SVZ, multipolar neurons move slowly tangentially, a process which may be required to explore the extracellular environment for putative polarity-inducing cues (Dimidschstein et al., 2013; Jossin, 2020). Next, multipolar neurons switch back to a bipolar state with the ventricle-oriented process developing into the axon. Bipolar projection neurons reattach to the radial glial fiber, migrate through the intermediate zone (IZ) and enter the cortical plate (CP) by using a locomotion mode (Hatanaka et al., 2004; Kriegstein and Noctor, 2004). Once the locomoting neurons reach the most superficial layer they detach from the radial glial fiber and perform terminal somal translocation to settle in their target position (Franco et al., 2011; Sekine et al., 2011, 2012).

In the last decades a large collection of signaling pathways including extracellular cues/receptors, cell adhesion molecules and their receptors, endocytic/exocytic regulators, intracellular signal transduction machinery, and transcription factors have been implicated in controlling the discrete sequential steps of radial projection neuron migration in healthy and disease conditions (Ayala et al., 2007; Buchsbaum and Cappello, 2019; Evsyukova et al., 2013; Hansen et al., 2017; Heng et al., 2010; Hippenmeyer, 2014; Kwan et al., 2012; Reiner et al., 2021). Thus far, most studies using loss-of-function paradigms focused almost exclusively on the cell-autonomous role of the genes encoding the above regulatory cues. Yet, there is accumulating evidence that non-cell-autonomous tissue-wide properties could substantially impact and/or contribute to the regulation of radial projection neuron migration (van den Berghe et al., 2014; Franco et al., 2011; Gorelik et al., 2017; Greenman et al., 2015; Hammond, 2004; Hammond et al., 2001; Hansen and Hippenmeyer, 2020; Hippenmeyer, 2014; Hippenmeyer et al., 2010; Nakagawa et al., 2019; Sanada et al., 2004; Yang et al., 2002; Youn et al., 2009) However, the nature and dynamics of such global tissue-wide effects remain unclear.

In our study we addressed this issue in a quantitative manner at single cell level and set out to establish experimental genetic paradigms enabling the separation of intrinsic cell-autonomous gene function from the contribution of tissue-wide effects. To this end we utilized Mosaic Analysis with Double Markers (MADM) technology (Contreras et al., 2021; Zong et al., 2005) which provides a quantitative platform *in situ* to 1) monitor neuronal migration with exquisite single cell resolution; and 2) probe the role of the genetic landscape and cellular environment on individual migrating neurons in a defined genetic context. Based on results from holistic single cell phenotypic analysis upon loss of gene function, we provide evidence that global tissue-wide effects predominate the overall cell-autonomous phenotype of migrating projection neurons.

## RESULTS

### MADM-based platform to probe cell-autonomous candidate gene function and the contribution of global tissue-wide effects at single cell level

In order to establish a quantitative assay to dissect cell-autonomous gene function and to measure the contribution of global tissue-wide cues/properties with single cell resolution we conceived a platform consisting of three genetic MADM-based paradigms: 1) Control-MADM; 2) *GeneX*-MADM (Mosaic-MADM); and 3) KO/cKO-*GeneX*-MADM (KO/cKO-MADM) (Figures 1A-1D and Figure S1). In Control-MADM, mice carrying MADM cassettes on a particular chromosome were combined with *Emx1*-Cre driver to generate experimental MADM mice with sparse fluorescent labeling in cortical projection neurons. Since these Control-MADM mice did not carry any loss-of-function (LOF) allele all cells were considered as ‘Control’ with some sparse cells expressing the fluorescent GFP and tdT markers enabling single cell tracing (Figure 1A). In *GeneX*-MADM with *Emx1*-Cre, a genetic mutation was coupled to one MADM cassette by meiotic recombination [see Materials and Methods (Amberg and Hippenmeyer, 2021; Contreras et al., 2021) for details] in a way that sparse homozygous mutant cells were always labeled in green (GFP^+^) and wild-type cells in red (tdT^+^), respectively (Figure 1B). In KO/cKO-*GeneX*-MADM with *Emx1*-Cre, the mutant allele of the gene of interest was coupled to both MADM cassettes resulting in full (KO) or conditional (cKO) tissue-specific knockout (*Emx1^+^* projection neuron lineage) with sparse fluorescent labeling in respective experimental MADM mice (Figure 1C). The key rationale in our overall assay was based on the following assumptions. First, in sparse genetic mosaic-MADM (where we ablated candidate gene function in just very few GFP^+^ migrating neurons) the observed phenotype of homozygous mutant cells implied the cell-autonomous gene function when compared to tdT^+^ Control cells. Second, the phenotype of individual mutant GFP^+^ cells in an all mutant environment (KO/cKO-MADM), of the same candidate gene as above, reflected the combination of 1) cell-autonomous loss of gene function; plus 2) global community effects in response to the loss of gene function in the vast majority of (if not all) cortical projection neurons across the entire tissue. Third, any quantitatively significant difference in the observed single cell mutant phenotype in mosaic-MADM when compared to KO/cKO-MADM indicated non-cell-autonomous and thus global tissue-wide effects (Figure 1D).

**Figure 1.**
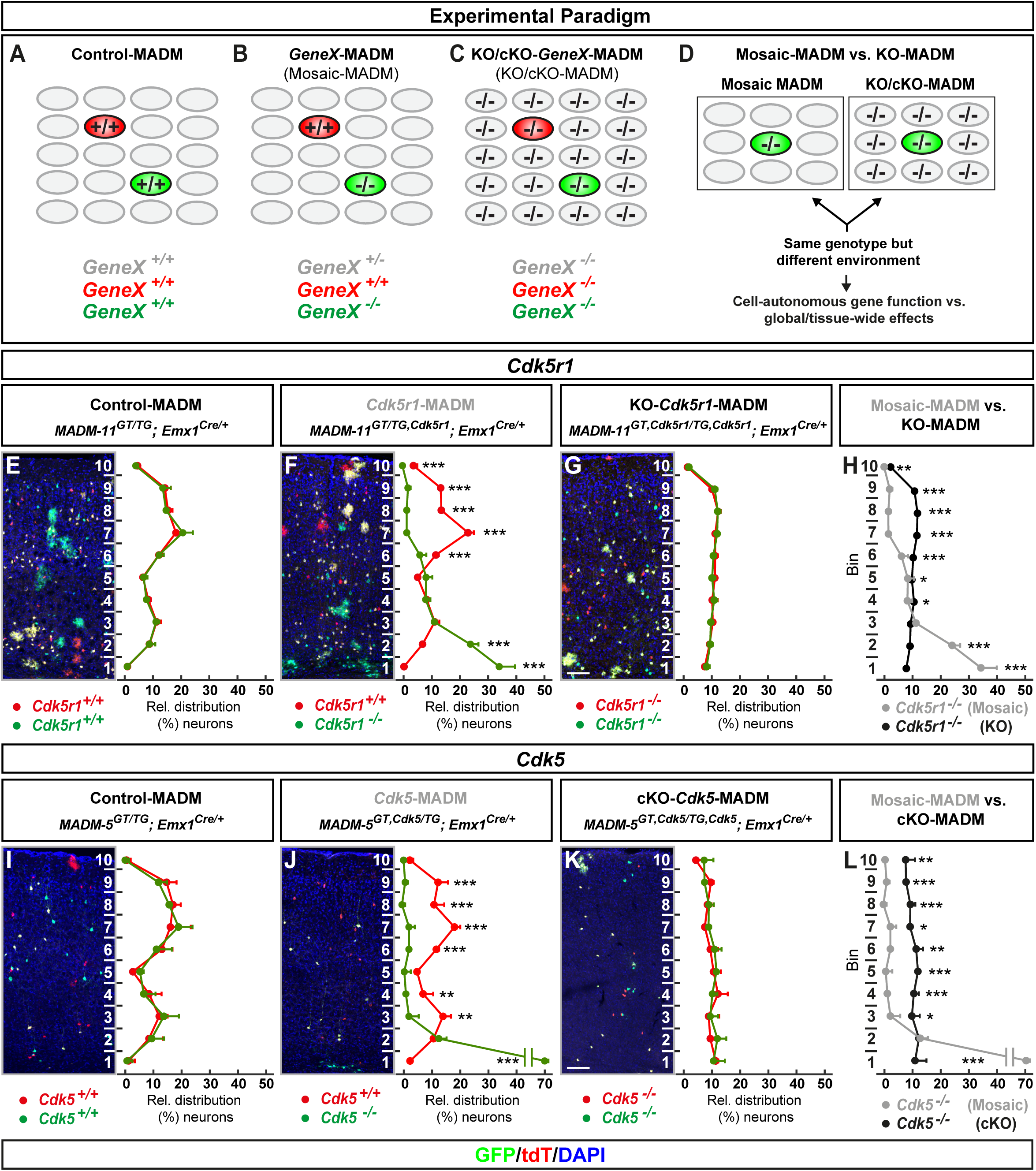
MADM analysis reveals that global tissue-wide effects predominate the cell-autonomous phenotype due to loss of p35/CDK5. (A-D) Experimental MADM paradigm to genetically dissect cell-autonomous gene function and non-cell-autonomous effects. (A) Control (Control-MADM: all cells *GeneX*^+/+^); (B) Sparse genetic mosaic (*GeneX*-MADM: only green cells are *GeneX*^-/-^ mutant, red cells are *GeneX*^+/+^ in an otherwise heterozygous *GeneX*^+/-^ environment); (C) Global/whole-tissue gene knockout (KO/cKO-*GeneX*-MADM: all cells mutant); (D) Direct phenotypic comparison of mutant cells in *GeneX*-MADM (Mosaic MADM) to mutant cells in KO/cKO-*Gene*-MADM (KO/cKO-MADM). Any significant difference in their respective phenotypes implies non-cell-autonomous effects. **(E-H)** Analysis of green (GFP^+^) and red (tdT^+^) MADM-labeled projection neurons in (E) Control-MADM (*MADM-11^GT/TG^;Emx1^Cre/+^*); (F) *Cdk5r1*-MADM (*MADM-11^GT/TG,Cdk5r1^;Emx1^Cre/+^*); and (G) KO-*Cdk5r1*-MADM (*MADM-11^GT,Cdk5r1/TG,Cdk5r1^;Emx1^Cre/+^*). Relative distribution (%) of MADM-labeled projection neurons is plotted in ten equal bins across the cortical wall. (H) Direct distribution comparison of *Cdk5r1^-/-^* mutant cells in *Cdk5r1*-MADM (grey) versus KO-*Cdk5r1*-MADM (black) distribution. **(I-L)** Analysis of green (GFP^+^) and red (tdT^+^) MADM-labeled projection neurons in (I) Control-MADM (*MADM-5^GT/TG^;Emx1^Cre/+^*); (J) *Cdk5*-MADM (*MADM-5^GT/TG,Cdk5^;Emx1^Cre/+^*); and (K) KO-*Cdk5*-MADM (*MADM-11^GT,Cdk5/TG,Cdk5^;Emx1^Cre/+^*). Relative distribution (%) of MADM-labeled projection neurons is plotted in ten equal bins (1-10) across the cortical wall. (L) Direct distribution comparison of *Cdk5^-/-^* mutant cells in *Cdk5*-MADM (grey) versus KO-*Cdk5*-MADM (black) distribution. Nuclei were stained using DAPI (blue). n=3 for each genotype with 10 (MADM-11) and 20 (MADM-5) hemispheres analyzed. Data indicate mean ± SD, *p<0.05, **p<0.01, and ***p<0.001. Scale bar: 100µm.

### Global tissue-wide effects predominate the cell-autonomous phenotype due to loss of p35/CDK5

The role of the p35/CDK5 signaling pathway in cortical projection neuron migration has been studied extensively (Delalle et al., 1997; Gupta et al., 2003; Hammond, 2004; Hatanaka et al., 2004; Ko et al., 2001; Ohshima, 2015). Thus with ample information available, based on LOF studies, this pathway provided an ideal case to dissect cell-autonomous requirement and the contribution of tissue-wide effects due to LOF. We first analyzed the single cell phenotype upon p35 (encoded by *Cdk5r1*, located on chr.11) - the main activator of CDK5 - ablation (Figure S2A). Therefore we generated Control-MADM (*MADM-11^GT/TG^;Emx1^Cre/+^*), *Cdk5r1*-MADM (*MADM-11^GT/TG,Cdk5r1^;Emx1^Cre/+^*), and KO-*Cdk5r1*-MADM (*MADM-11^GT,Cdk5r1/TG,Cdk5r1^;Emx1^Cre/+^*). Next we quantified the relative distribution of red and green MADM-labeled cortical projection neurons (terminal position) in somatosensory cortex at postnatal day (P) 21 to assess overall migration capacity in the above MADM paradigms (Figures 1E-1G). In Control-MADM red and green (both Control) projection neurons distributed similarly but in a defined fashion (relatively higher number in upper layers) across 10 bins as previously reported (Hippenmeyer et al., 2010) (Figure 1E). In *Cdk5r1*-MADM, the green homozygous *Cdk5r1*^-/-^ mutant neurons accumulated in lower layers, indicative of the cell-autonomous requirement for radial migration (Gupta et al., 2003; Hammond, 2004), while red Control neurons showed a distribution pattern similar to red/green cells in Control-MADM (Figure 1F). In contrast, red and green (both homozygous *Cdk5r1*^-/-^ mutant) neurons in KO-*Cdk5r1*-MADM showed similar pattern when compared to each other but clearly different distribution pattern when compared to Control and mutant cells in Control-MADM and/or *Cdk5r1*-MADM, respectively (Figure 1G). Strikingly, the distribution of green homozygous *Cdk5r1*^-/-^ mutant neurons in *Cdk5r1*-MADM was significantly distinct when compared to green homozygous *Cdk5r1*^-/-^ mutant neurons in KO-*Cdk5r1*-MADM despite that both cell populations had the same genotype (Figure 1H). Thus, the cell-autonomous phenotype (i.e. terminal location) of *Cdk5r1*^-/-^ mutant cortical projection neurons was significantly influenced by the genetic tissue-wide landscape.

To corroborate the above finding we established a second MADM platform by using MADM cassettes inserted in chr.5 (i.e. MADM-5) to more directly assay the consequences of *Cdk5* (located on chr.5) loss of function (Figure S2B). We thus generated Control-MADM (*MADM-5^GT/TG^;Emx1^Cre/+^*), *Cdk5*-MADM (*MADM-5^GT,Cdk5/TG^;Emx1^Cre/+^*) and cKO-*Cdk5*-MADM (*MADM-5^GT,Cdk5/TG,Cdk5^;Emx1^Cre/+^*); and analyzed the terminal distribution of red and green cells across the somatosensory cortex at P15 (since cKO-*Cdk5*-MADM animals tend to die soon thereafter). Again, in Control-MADM red and green cells (both wild-type) distributed similarly when compared to each other, and similarly when compared to red and green cells in Control-MADM using MADM-11 (Figures 1E and 1I). In mosaic *Cdk5*-MADM, green mutant *Cdk5*^-/-^ cells accumulated in lower layers and the white matter (Figure 1J). The migration phenotype of *Cdk5*^-/-^ cells in *Cdk5*-MADM appeared slightly stronger than the one of *Cdk5r1*^-/-^ cells in *Cdk5r1*-MADM (Figure 1F and J). The small phenotypic difference could presumably be due to possible compensation by p39 upon loss of p35 and thus some minor residual CDK5 activity in *Cdk5r1*^-/-^ cells (Ko et al., 2001; Su and Tsai, 2011). In any case, red and green (both homozygous *Cdk5*^-/-^ mutant) neurons in KO-*Cdk5*-MADM showed clearly different distribution pattern when compared to Control and mutant cells in Control-MADM and/or *Cdk5*-MADM, respectively (Figure 1I, J and K). Again, the distribution of green, homozygous *Cdk5*^-/-^ mutant neurons, in *Cdk5*-MADM was significantly distinct when compared to green homozygous *Cdk5*^-/-^ mutant neurons in KO-*Cdk5*-MADM despite that both cell populations had the same genotype (Figure 1L). Altogether, our data indicate that the state of the genetic environment, surrounding cells carrying mutations in genes regulating radial projection neuron migration in the developing neocortex, contributes critically to the overall phenotype of individual cells.

### Developmental progression of tissue-wide effects impacting phenotypic manifestation upon sparse and global KO of p35/CDK5

To determine the emergence of non-cell-autonomous tissue-wide effects influencing radial neuron migration we pursued developmental time course analysis. We utilized the same MADM-based paradigms as described in the above section to visualize the single cell phenotype upon sparse and global elimination of *Cdk5r1* (Figures 2A-2L) and *Cdk5* (Figures 2M-2X), respectively. At embryonic day (E) 14 no phenotypic difference - i.e. relative vertical distribution of mutant cells in mosaic versus cKO/KO - in VZ/SVZ and IZ was observed. However, small but significant differences were apparent in the upper bins, corresponding to the emerging CP, in both *Cdk5r1* and *Cdk5* comparative sparse and global MADM deletion paradigms (Figures 2D and 2P). At E16 (Figures 2E-2F and 2Q-2R) and P0 (Figures 2I-2J and 2U-2V) time points, based on sparse mosaic MADM paradigms, the cell-autonomous phenotypes demonstrated critical requirement for *Cdk5r1/Cdk5* in migrating projection neurons to enter the developing cortical plate. In effect, although mutant cells could migrate from the VZ/SVZ through IZ, *Cdk5r1^-/-^* and *Cdk5^-/-^* mutant cells accumulated below the CP. The phenotypic manifestation became progressively stronger as the development of the CP proceeded. In contrast, mutant cells upon global *Cdk5r1/Cdk5* ablation in KO/cKO appeared to distribute more evenly across the developing cortical wall (Figures 2G, 2K, 2S and 2W). Thus, the emerging cell-autonomous phenotype upon *Cdk5r1/Cdk5* ablation (deficit to enter the developing CP) was significantly affected by the global genetic constitution of the developing cortical wall.

**Figure 2.**
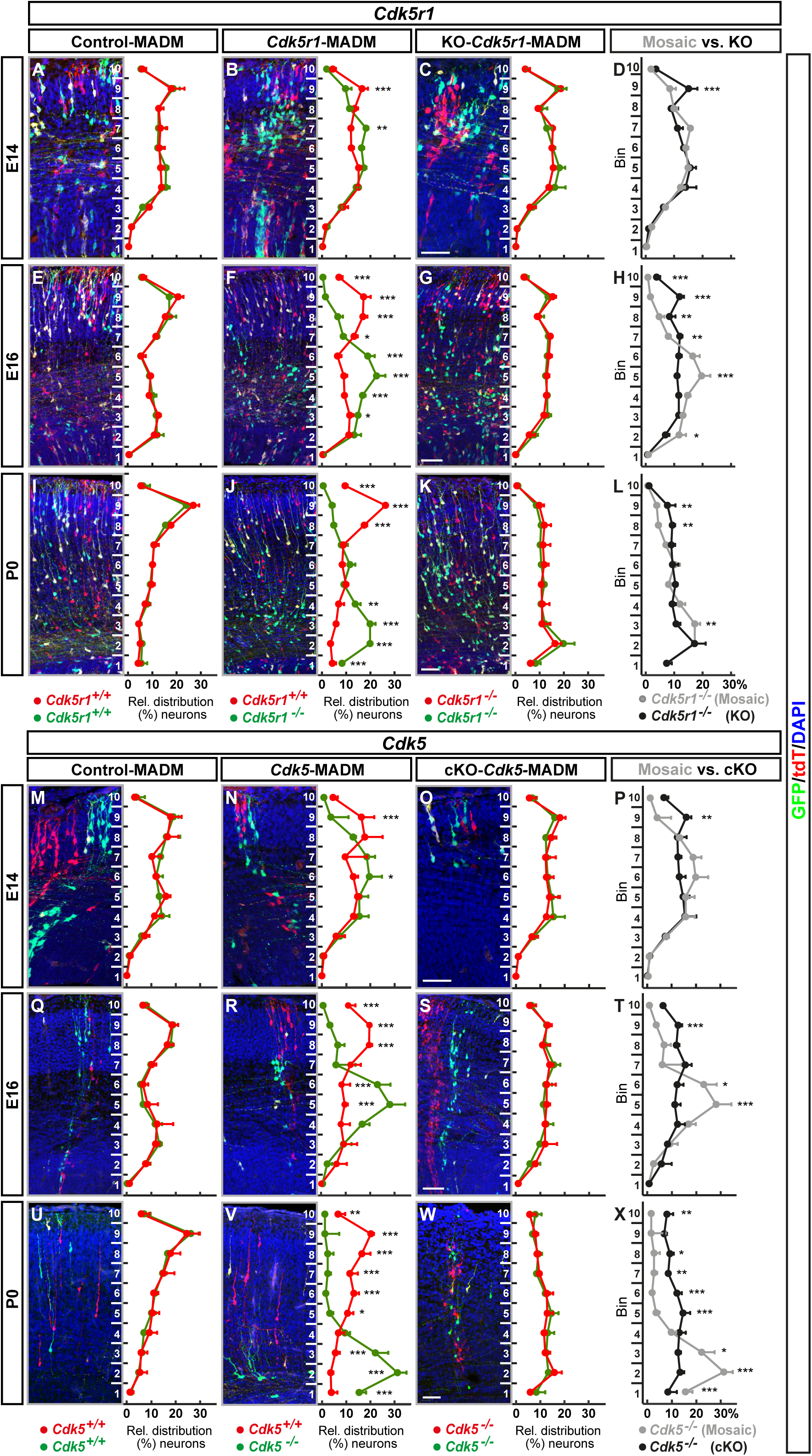
Developmental time course analysis of MADM-based sparse and global KO of p35/CDK5. (A-L) Analysis of green (GFP^+^) and red (tdT^+^) MADM-labeled projection neurons in (A, E, I) Control- MADM (*MADM-11^GT/TG^;Emx1^Cre/+^*); (B, F, J) *Cdk5r1*-MADM (*MADM-11^GT/TG,Cdk5r1^;Emx1^Cre/+^*); and (C, G, K) KO-*Cdk5r1*-MADM (*MADM-11^GT,Cdk5r1/TG,Cdk5r1^; Emx1^Cre/+^*) at E14 (A-D), E16 (E-H), and P0 (I-L). Relative distribution (%) of MADM-labeled projection neurons is plotted in ten equal bins across the developing cortical wall. (D, H, L) Direct distribution comparison of *Cdk5r1^-/-^* mutant cells at E14 (D), E16 (H), and P0 (L) in *Cdk5r1*-MADM (grey) versus KO-*Cdk5r1*-MADM (black) distribution. **(M-X)** Analysis of green (GFP^+^) and red (tdT^+^) MADM-labeled projection neurons in (M, Q, U) Control-MADM (*MADM-11^GT/TG^;Emx1^Cre/+^*); (N, R, V) *Cdk5*-MADM (*MADM-11^GT/TG,Cdk5^;Emx1^Cre/+^*); and (O, S, W) KO-*Cdk5*-MADM (*MADM-11^GT,Cdk5/TG,Cdk5^; Emx1^Cre/+^*) at E14 (M-P), E16 (Q-T), and P0 (U-X). Relative distribution (%) of MADM-labeled projection neurons is plotted in ten equal bins across the developing cortical wall. (P, T, X) Direct distribution comparison of *Cdk5^-/-^* mutant cells at E14 (P), E16 (T), and P0 (X) in *Cdk5*-MADM (grey) versus KO-*Cdk5*-MADM (black) distribution. Nuclei were stained using DAPI (blue). n=3 for each genotype with 10 (MADM-11) and 20 (MADM-5) hemispheres analyzed. Data indicate mean ± SD, *p<0.05, **p<0.01, and ***p<0.001. Scale bars: 100µm.

### Distinct projection neuron migration dynamics upon sparse and global KO of p35/CDK5

To directly assess cortical projection neuron migration dynamics we measured physical movement *in-situ* by time-lapse imaging (Figure 3A). We analyzed neuronal migration in embryonic brain slices at E16 when cortical projection neurons can be found at all stages in their sequential radial migration trajectory. We capitalized upon the exquisite single cell resolution in the three - Control-MADM (Figure 3B, Movie S1), *Cdk5r1*-MADM (Figure 3C, Movie S2), and KO-*Cdk5r1*-MADM (Figure 3D, Movie S3) - experimental paradigms and recorded confocal images at 15min intervals over an extended period of >15 hours. Frames were generated from individual confocal stacks and processed for analysis (see Materials and Methods for details). From all imaging data we could systematically analyze migration dynamics in movies of at least 12 hours (725min) for all three genetic MADM paradigms. We divided the developing cortical wall into two (lower and upper) bins. The lower bin comprised the VZ/SVZ and IZ whereas the upper bin corresponded to the emerging CP. To empower our analysis we employed a semi-automated tracking method to eliminate any experimenter bias (see Materials and Methods). We focused the analysis on the following four populations of MADM-labelled cells: Control (red and green neurons in Control-MADM), Mosaic-control (red Control neurons in *Cdk5r1*-MADM), Mosaic-mutant (green *Cdk5r1*^-/-^ mutant neurons in *Cdk5r1*-MADM), and KO-mutant (red and green *Cdk5r1*^-/-^ mutant neurons in KO-*Cdk5r1*-MADM). First, we evaluated the relative distribution of cells with distinct genotypes across the two (upper and lower) bins at the start (t=0) and the end (t=725min) of the recorded time series (Figures 3E and 3F). We noticed that at the start time point roughly 50% of Control, Mosaic-control and KO-mutant (*Cdk5r1*^-/-^) cells were located in the upper and ∼50% in the lower bin, respectively. In contrast, Mosaic-mutant (*Cdk5r1*^-/-^) cells in *Cdk5r1*-MADM failed to distribute evenly between upper and lower bins, and accumulated significantly in the lower bin with much less cells in the upper bin. These data demonstrate that the cell-autonomous *Cdk5r1* function, to promote the transition from the IZ into the CP (e.g. Figures 2B, 2F and 2J), is preserved in our *in vitro* migration assay. At the end time point there was a trend whereby Control and Mosaic-control were slightly overrepresented in the upper bin relative to the amount of cells in the lower bin. However, Mosaic-mutant neurons still displayed strong bias towards the lower bin, and KO-mutant cells remained at about 50/50 distribution at the end versus start time point.

**Figure 3.**
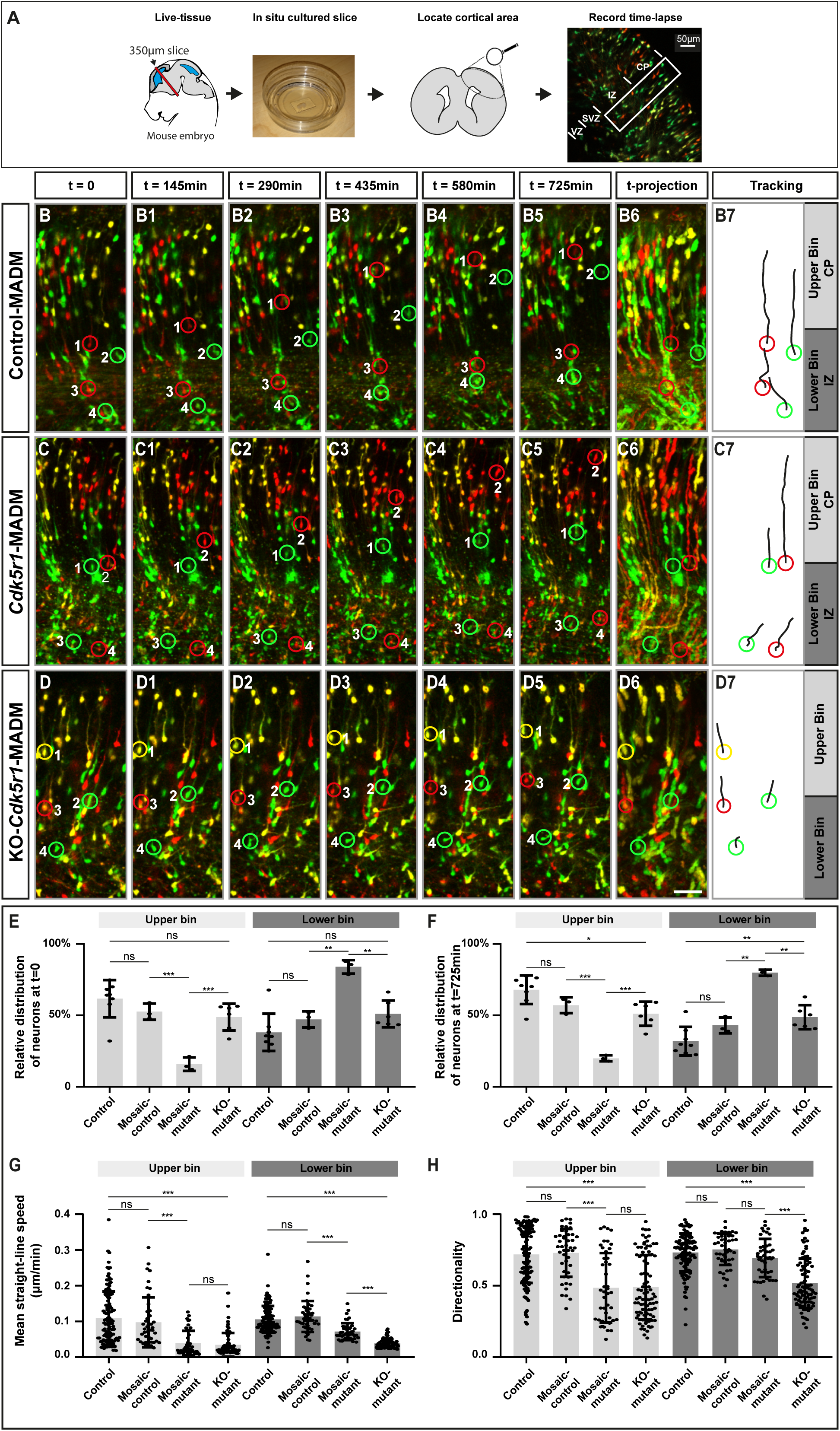
Projection neuron migration dynamics upon sparse and global KO of p35/CDK5. **(A)** Experimental setup for time-lapse imaging of MADM-labeled neurons in developing somatosensory cortex at E16. **(B-D)** Time-lapse imaging of (B-B5) Control-MADM (*MADM-11^GT/TG^;Emx1^Cre/+^*), (C-C5) *Cdk5r1*-MADM (*MADM-11^GT/TG,Cdk5r1^;Emx1^Cre/+^*), and (D-D5) KO-*Cdk5r1*-MADM (*MADM-11^GT,Cdk5r1/TG,Cdk5r1^; Emx1^Cre/+^*). (B6, C6, D6) 12h time-projection of sequential images in IZ (Lower Bin) and emerging CP (Upper Bin) at 15min framerate. (B7, C7, D7) Tracking trajectories of indicated neurons (red and green rings) in Control-MADM (B7), *Cdk5r1*-MADM (C7), and KO-*Cdk5r1*-MADM (D7). **(E)** Relative distribution of neurons at the start of the time-lapse t=0 for each replicate time-lapse per genotype. **(F)** Relative distribution of neurons at the end of the time-lapse t=725min for each replicate time-lapse per genotype. **(G)** Mean straight-line speed of the top 15 tracks per replicate time-lapse per genotype. **(H)** Directionality of the top 15 tracks per replicate time-lapse per genotype. n=3 videos from >2 independent animals. Data indicate mean ± SD, *p<0.05, **p<0.01, and ***p<0.001. Scale bar: 40μm.

Next we assessed migration speed (Figure 3G) and directionality (Figure 3H). Control and Mosaic-control neurons did migrate at equal speeds in the upper and lower bins while Mosaic-mutant and KO-mutant migrated at significantly slower speed in both compartments. Interestingly, KO-mutant neurons migrated even significantly slower than Mosaic-mutant neurons in the lower bin. While Control and Mosaic-control neurons showed highly directional (vertical orientation from VZ toward pial surface) migration behavior in both, the upper and lower bins, Mosaic-mutant and KO-mutant cells in the upper bin showed significantly less directional movement. In the lower bin, Mosaic-mutant cells were not affected in directional movement when compared to Control cells but migrating KO-mutant neurons showed significantly less directionality. Altogether, KO-mutant neurons migrated significantly slower and showed significantly less directional movement in the lower bin when compared to Mosaic-mutant neurons. Thus, while both populations were *Cdk5r1*^-/-^, their environments were distinct (i.e. *Cdk5r1*^-/-^ only in KO-mutant). We therefore conclude that dynamics and directionality of radially migrating cortical projection neurons is critical dependent on the genetic landscape of the cellular environment.

### *In silico* modelling reveals cell-autonomous force generation and directionality in concert with environmental tissue resistance as minimal parameters instructing cortical projection neuron migration

The above data clearly demonstrated that the interplay of cell-autonomous gene function with non-cell-autonomous tissue-wide properties controls overall efficiency of cortical projection neuron migration. In order to obtain a more quantitative model we set out to define a minimal set of physical parameters sufficient to encompass migration dynamics on a statistical level. We based our approach on the cell tracking data which we extracted from the time-lapse imaging experiments described above. We first analyzed the trajectories of Control cells from Control-MADM (Figure 4A), KO-mutant cells from KO-*Cdk5r1*-MADM (Figure 4B), and Mosaic-control and Mosaic-mutant cells from *Cdk5r1*-MADM (Figure 4C); and plotted the overall experimental velocity distribution of each cell population as a reference (Figure 4D). More specifically, the temporal change in position resulted in a distribution of velocities (i.e. normalized velocity) throughout the developing cortical wall ranging from the ventricle to the pia (y axis in Figure 4D). For Control, Mosaic-control and to some extent Mosaic-mutant (cells in CP considered as ‘escapers’ being able to cross the IZ/CP border despite loss of *Cdk5r1*) neurons the distribution showed strong velocity peak within the lower half of the tissue (lower bin) followed by a sharp decrease in velocity around the border of the upper bin. However, KO-mutant cells did not display such characteristics but rather showed a more or less even distribution throughout the tissue. The velocity distributions and thus migration dynamics at the IZ-CP (lower-upper bin) border with a positional change in velocity implied a change in tissue architecture and suggested a difference in ‘stiffness’ between the two compartments. Indeed, atomic force microscopy measurements (i.e. Young’s moduli *E* as proxy for stiffness or resistance) previously indicated distinct stiffness of individual cortical compartments at different developmental stages (Iwashita et al., 2014).

**Figure 4.**
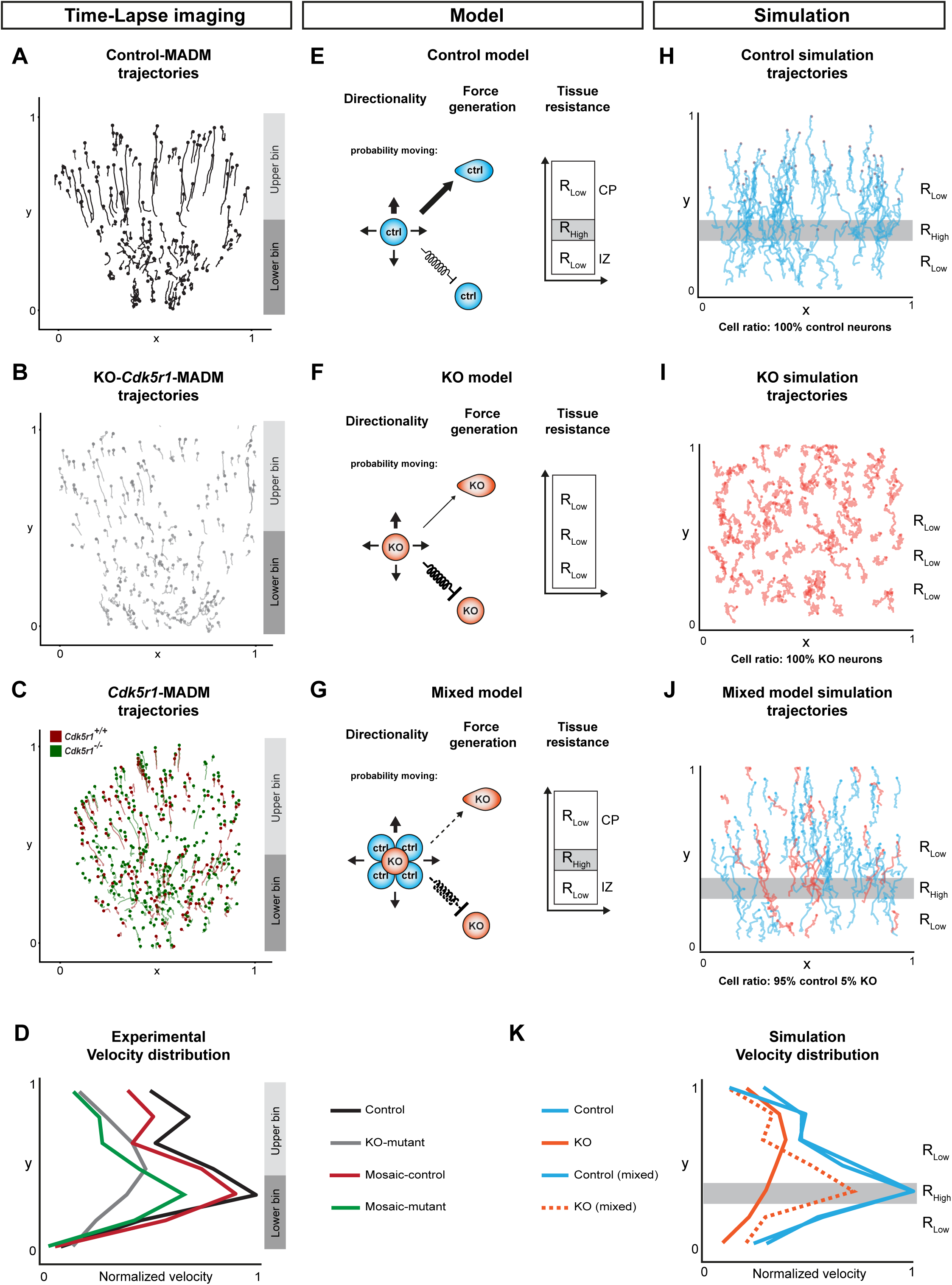
*In silico* modelling of neuronal migration dynamics upon MADM-based *Cdk5r1* ablation. **(A-C)** Representative migration trajectories in (A) control-MADM, (B) KO-*Cdk5r1*-MADM, and (C) *Cdk5r1*-MADM. **(D)** Normalized velocity distributions from experimental data in control-MADM (n=4), KO-*Cdk5r1*-MADM (n=3), and *Cdk5r1*-MADM (n=3). **(E)** Model of neuron migration in Control environment with corresponding resistance zones. The thickness of the arrows indicates the probability to move in any direction. The directionality bias is defined as 65% in pial-direction for Control. **(F)** Model of neuron migration in global *Cdk5r1* KO environment with a single resistance zone. The thickness of the arrows indicates the probability to move in any direction, here the directionality bias is defined as 51% in pial-direction. **(G)** Mixed model indicating cross interactions of *Cdk5r1^-/-^* mutant and Control cells, respectively. The directionality bias is defined in pial-direction as a function of N_Ctrl/N_Mut ratio min=51%, max=65%. The thickness of the arrows indicates the probability of the mutant neuron to move in any direction. **(H-I)** Simulation of migration trajectories in (H) Control model, (I) global *Cdk5r1* KO model, and (J) mixed model (95% Ctrl, 5% KO). Resistant zones are indicated accordingly. **(K)** Normalized velocity distributions of simulation trajectories.

Next, from the experimental data we extracted two cell-intrinsic parameters, directionality (*ρ*) and force generation (*α*), and inferred one extrinsic parameter defining the environmental tissue resistance (*R_i_*) (Figure 4E, see also Materials and Methods). For the latter, in a reductionist model we assumed that IZ and CP are relatively homogenous and that overall stiffness can be attributed to differences in physical pore-size distribution. In other words, when a cell migrates through a porous environment it will have to squeeze through different pores in order to advance. Thus the overall resistance that the cells are experiencing decreases with increasing pore size. The magnitude of the resistance fields *R_i_* is defined as a free parameter of our system. As described above, in Control conditions migrating cells at the IZ-CP border seem to struggle in order to squeeze through. In contrast, due to virtual inexistence of physiological layer structure in KO-*Cdk5r1*-MADM the migration dynamics appear independent of the location and thus of the *R_tissue_*. To integrate the two described environments in the model, we designed an environment with three compartments and two different resistances (*R_low_* and *R_high_*). In the control environment the layers were defined by *R_Control,tissue_*=*R_1_*,*R_2_*,*R_3_* for *R_2_*>*R_1_*∼*R_3_*, with *R_2_* roughly corresponding to the IZ-CP border (Figure 4E and Table S2) while in KO-*Cdk5r1*-MADM environment we set *R_KO,tissue_*=*R_KO_* with magnitude on the order of *R_1_*,*R_3_* (Figure 4F and Table S2). The tissue resistance parameters in combination with directional bias *ρ* and force generation *α* allowed us to model cells as persistent random walkers (Figure S3A, see also Materials and Methods for details).

Based on our model, we next simulated Control neuron migration (Figure 4H) and could very closely mimic the experimentally observed migration behavior and velocity distribution (Figures 4A, 4D, 4H, and 4K). Likewise, the simulation of KO-mutant cells (Figure 4I) in a uniformly low resistance environment *R_KO,tissue_* [with reduced directionality parameter *ρ* and force scaling parameter *α*, respectively] matched our observations in experimental data from KO-*Cdk5r1*-MADM paradigm (Figure 4B, 4D, 4I, and 4K).

Next we utilized a ‘mixed model’ where we introduced KO-mutant cells into a surrounding of Control cells with according control tissue resistance environment (Figure 4G). Here we introduced a linear coupling of directionality *ρ* and force generation coefficient *α* with the ratio of Control to KO-mutant cells (Figure 4J, Figure S3B). The simulation conducted with the mixed model mimicked the dynamics and distributions we detected in *Cdk5r1*-MADM where mutant neurons appeared more dynamic than in KO-*Cdk5r1*-MADM (Figures 4J-4K). Further we could model the emergence of the non-cell-autonomous effects with increasing ratio of Control to KO-mutant cells (Figure S3C). The more KO-mutant cells present in the simulation, the more uniform the velocity distribution appeared across the vertical axis of the cortical wall. At a ratio of 4/96% (KO-mutant to Control), the KO-mutant cells showed velocity distribution like Mosaic-mutant cells and Control cells (similar to the observation in the experimental data). In contrast, at a ratio 10/90%, the KO-mutant cells displayed velocity distribution as observed in KO-*Cdk5r1*-MADM.

Moreover, in our model we found that the transition between the two distributions appeared abruptly between 5% and 6% abundance of KO-mutant cells. In summary, our *in silico* model can provide an accurate quantitative replicate of the migration behavior and dynamics as measured *in situ*. Our data further demonstrate that the relative amount of *Cdk5r1^-/-^* mutant cells and thus the overall genetic landscape within the cortical tissue critically impacts on the migration dynamics of individual cells.

### Distinct deregulation of gene expression in *Cdk5r1^-/-^* mutant cells upon sparse and global KO

Our data so far indicates distinct single cell *in vivo* phenotypes in *Cdk5r1^-/-^* mutant cells depending on the genetic state of the environment. By using *in silico* modeling we also showed that the tissue resistance (in combination with cell-intrinsic directionality and force generation capability) along the migration path represented a most critical physical parameter. To obtain a hint on molecular correlates mirroring the distinct observed phenotypes we next pursued transcriptome analysis. We specifically isolated green GFP^+^ MADM-labeled cortical projection neurons by FACS (Laukoter et al., 2020a, 2020b) in Control-MADM (*Cdk5r1^+/+^*), *Cdk5r1*-MADM (*Cdk5r1^-/-^*) and KO-*Cdk5r1*-MADM (*Cdk5r1^-/-^*) at E13, E16 and P0, respectively (Figure 5A). We performed RNA-sequencing of small bulk samples using SMARTer technology, followed by bioinformatics analysis (see Materials and Methods). First we analyzed *Cdk5r1* expression and found that in both (sparse and global KO) deletion paradigms, the *Cdk5r1* expression level was close to zero while in Control (*Cdk5r1^+/+^*) cells substantial *Cdk5r1* expression was evident (Figure S4). These data validated the MADM-based *Cdk5r1* ablation paradigms. We next analyzed differentially expressed genes (DEGs) in *Cdk5r1*-MADM and KO-*Cdk5r1*-MADM in comparison to Control-MADM (Figure 5B). At P0 we identified 3 significant DEGs in *Cdk5r1*-MADM which was in stark contrast to the KO-*Cdk5r1*-MADM cells where we identified 1056 DEGs (padj<0.05, DESeq2). Thus depending on the state of the genetic environment (global *Cdk5r1* KO or not), three orders of magnitude higher number of DEGs was observed in individual *Cdk5r1^-/-^* mutant cells. Next we directly compared the difference in DEGs between *Cdk5r1^-/-^* mutant cells in sparse versus global KO. We observed a progressive increase of DEGs during development with slightly more genes showing downregulation (Figure 5C-5D). To obtain more insight into the biological DEG functions we performed gene ontology (GO) enrichment analysis at P0 on the 670 up- and 927 downregulated genes (Figure 5E). The top enriched GO terms for the downregulated genes (blue) were highly significant (padj<2.45x10^-9^, hypergeometric test) and associated with extracellular matrix (ECM), cell membrane, and cell adhesion. GO term enrichment in the upregulated (yellow) genes was not significant (padj>0.5) and associated with synaptic terms (Figure 5E). In summary, transcriptome analysis identified downregulation of ECM, membrane associated and cell adhesion genes as major classes of DEGs in *Cdk5r1^-/-^* cells in global KO-*Cdk5r1*-MADM but not sparse mosaic *Cdk5r1*-MADM.

**Figure 5.**
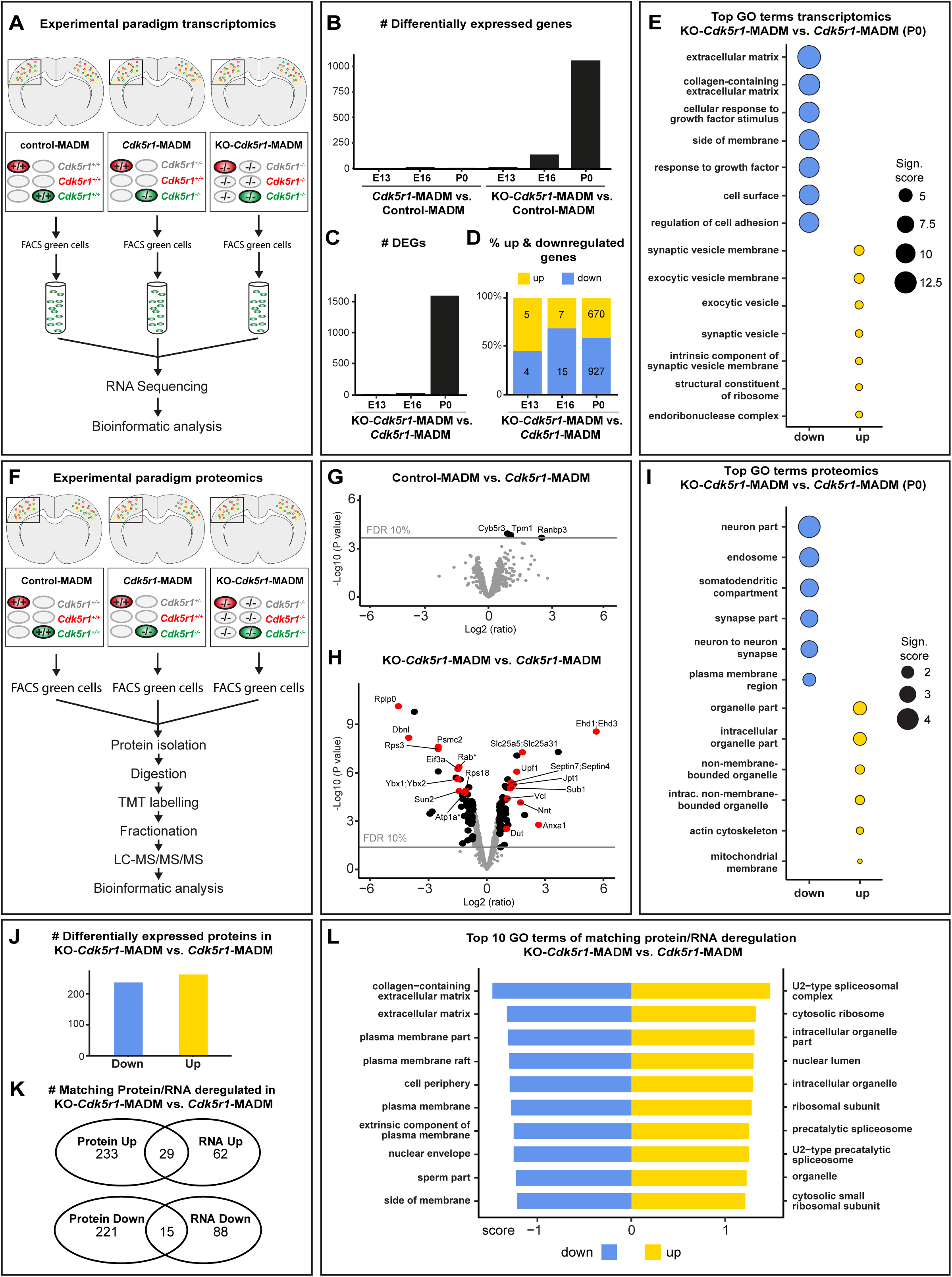
Gene and protein expression in *Cdk5r1^-/-^* mutant cells upon sparse and global KO. **(A)** Experimental paradigm and pipelines for gene expression profiling in Control-MADM (left), *Cdk5r1*- MADM (middle), and KO-*Cdk5r1*-MADM (right). **(B)** Number of differentially expressed genes (DEGs) in *Cdk5r1*-MADM and KO*-Cdk5r1*-MADM versus Control at E13, E16, and P0. **(C)** Number of DEGs in KO*-Cdk5r1*-MADM versus *Cdk5r1-*MADM at E13, E16, and P0. **(D)** Percentage of up- and downregulated genes in KO*-Cdk5r1*-MADM versus *Cdk5r1-*MADM at E13, E16, and P0. **(E)** Top GO terms associated with genes in (C and D) at P0. Note that GO term enrichments for upregulated genes are non-significant. **(F)** Experimental paradigm and pipelines for proteome profiling in Control-MADM (left), *Cdk5r1*-MADM (middle), and KO-*Cdk5r1*-MADM (right). **(G)** Volcano plot showing deregulated proteins in Control-MADM versus *Cdk5r1*-MADM comparison at P0. Note that only 3 proteins were significantly upregulated. **(H)** Volcano plot showing deregulated proteins KO*-Cdk5r1*-MADM versus *Cdk5r1*-MADM comparison at P0. **(I)** Top enriched GO terms associated with genes encoding the proteins as shown in (H). **(J)** Number of genes associated with differentially expressed proteins in KO-*Cdk5r1*-MADM versus *Cdk5r1*-MADM. Note that criteria for significant differential expression were relaxed compared to (H). **(K)** Venn diagrams indicating the overlap of deregulated genes in transcriptomic and proteomic datasets in KO- *Cdk5r1*-MADM versus *Cdk5r1*-MADM. **(L)** Top 10 GO-terms associated with gene sets that are up- and downregulated in both (transcriptomic and proteomic) data sets (overlap in K).

### Deregulation of ECM and cell adhesion proteins upon global but not sparse *Cdk5r1^-/-^* KO

Next we determined putative differences in *Cdk5r1^-/-^* mutant cells upon global versus sparse KO at the proteomic level. We thus isolated the green GFP^+^ MADM-labeled cortical projection neurons by FACS (Laukoter et al., 2020a, 2020b) in Control-MADM (*Cdk5r1^+/+^*), *Cdk5r1*-MADM (*Cdk5r1^-/-^*) and KO- *Cdk5r1*-MADM (*Cdk5r1^-/-^*) (Figure 5F), followed by liquid chromatography-tandem mass spectrometry (LC-MS/MS) (Figure 5F, see also Materials and Methods). First we compared the proteome of *Cdk5r1*-MADM mutant neurons to Control-MADM cells whereby we identified (only) three significantly deregulated proteins (Figure 5G). However, when mutant cells in *Cdk5r1*-MADM (*Cdk5r1^-/-^*) were compared with mutant cells in KO-*Cdk5r1*-MADM (*Cdk5r1^-/-^*), we identified 59 and 61 significantly up- and downregulated proteins, respectively (Figure 5H). GO-term enrichment analysis (of the genes associated with the above differentially expressed proteins) indicated top GO terms associated with membrane and neuron-neuron contact among the downregulated group, thus corroborating our findings from the transcriptome analysis. GO-terms for the upregulated group were mainly associated with intracellular processes such as organelles (Figure 5I). Having proteomic and transcriptomic information at hand we next analyzed putative overlap in the two data sets. Since the direct correlation between transcriptome and proteome is complex we lowered the thresholds for differential expression (see Materials and Methods). We focused the analysis on 1060 gene annotations, which were informative in both data sets, and identified 262 downregulated and 236 upregulated DEGs from the proteomics data set (Figure 5J). Strikingly, 29 upregulated and 15 downregulated genes were common to both transcriptomic and proteomic analyses (Figure 5K). GO-term enrichment analysis identified ECM and membrane-associated GO terms among the down regulated gene group and GO terms related to cell-intrinsic entities among the upregulated gene group (Figure 5L). In summary, the above analysis revealed significant downregulation of mRNAs and proteins related to membrane, cell adhesion and ECM, specifically in *Cdk5r1^-/-^* mutant cells upon global tissue-wide but not sparse KO.

### Global tissue-wide effects predominate the cell-autonomous phenotype due to loss of *Dab1*

Global elimination of p35/CDK5 signaling in cortical projection neurons results in dominant tissue-wide effects that affect the individual cell-autonomous phenotype. To evaluate whether such observation is specific to p35/CDK5 signaling pathway we next investigated the consequences of ablation of Reelin/DAB1 signaling [which operates in parallel to CDK5 (Bock and May, 2016; Keshvara et al., 2002; Kwon and Tsai, 1998; Ohshima, 2015)]. We therefore generated MADM-based *Dab1* (located on chr.4) ablation paradigms using MADM-4 platform (Figure S2C). Next we analyzed radial neuron migration in Control-MADM (*MADM-4^GT/TG^;Emx1^Cre/+^*), *Dab1*-MADM (*MADM-4^GT/TG,Dab1^;Emx1^Cre/+^*), and KO-*Dab1*- MADM (*MADM-4^GT,Dab1/TG,Dab1^;Emx1^Cre/+^*) at E16, P0, and P21 in somatosensory cortex (Figures 6A-D and S5). In *Dab1*-MADM the majority of *Dab1^-/-^* mutant neurons was positioned in a biased manner, reflecting the cell-autonomous *Dab1* function (Franco et al., 2011; Hammond et al., 2001; Sanada et al., 2004) within lower bins in the distribution charts (i.e. below CP during earlier development and lower cortical layers at later stages). Conversely, *Dab1^-/-^* mutant neurons in KO-*Dab1*-MADM showed much more even distribution throughout the cortical wall (Figures 6A-D and S5).

**Figure 6.**
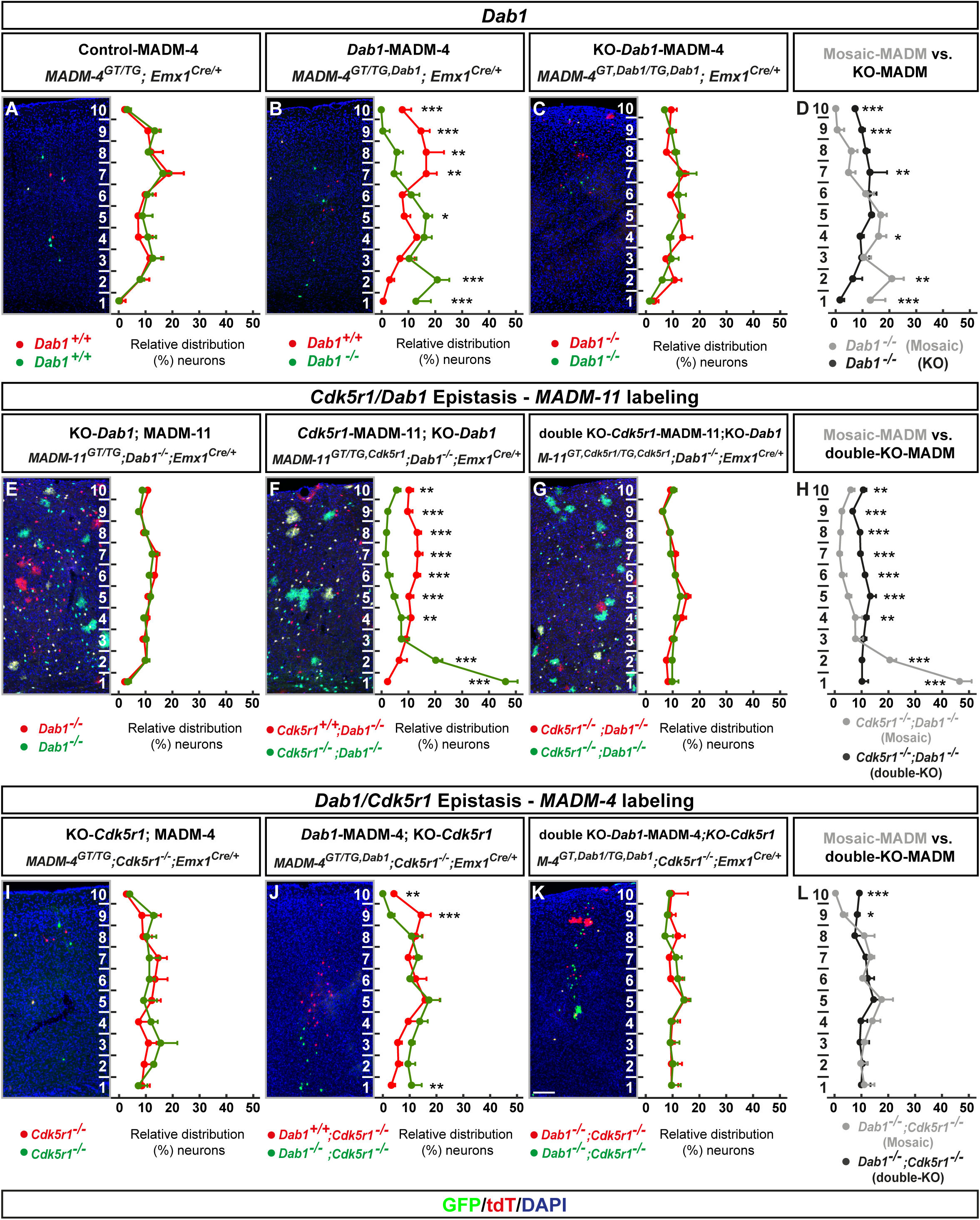
MADM-based analysis of *Dab1* and *Cdk5r1*/*Dab1* epistasis. **(A-D)** Analysis of green (GFP^+^) and red (tdT^+^) MADM-labeled projection neurons in (A) Control-MADM (*MADM-4^GT/TG^;Emx1^Cre/+^*); (F) *Dab1*-MADM (*MADM-4^GT/TG,Dab1^;Emx1^Cre/+^*); and (G) KO-*Dab1*-MADM (*MADM-4^GT,Dab1/TG,Dab1^;Emx1^Cre/+^*). Relative distribution (%) of MADM-labeled projection neurons is plotted in ten equal bins across the cortical wall. (H) Direct distribution comparison of *Dab1^-/-^* mutant cells in *Dab1*-MADM (grey) versus KO-*Dab1*-MADM (black) distribution. **(E-H)** *Cdk5r1*/*Dab1* epistasis in MADM-11 labeling background. Analysis of green (GFP^+^) and red (tdT^+^) MADM-labeled projection neurons in (E) KO-*Dab1*; MADM-11 (*MADM-11^GT/TG^;Dab1^-/-^;Emx1^Cre/+^*), (F) *Cdk5r1*-MADM-11;KO-*Dab1* (*MADM-11^GT/TG,Cdk5r1^;Dab1^-/-^;Emx1^Cre/+^*), and (G) double-KO-*Cdk5r1*-MADM-11;KO-*Dab1* (*MADM-11^GT,Cdk5r1/TG,Cdk5r1^;Dab1^-/-^;Emx1^Cre/+^*). Relative distribution (%) of MADM-labeled projection neurons is plotted in ten equal bins across the cortical wall. (H) Direct distribution comparison of *Cdk5r1^-/-^* mutant cells upon sparse (grey) and global (black) KO in *Dab1^-/-^* mutant background. **(I-L)** *Dab1/Cdk5r1* epistasis in MADM-4 labeling background. Analysis of green (GFP^+^) and red (tdT^+^) MADM-labeled projection neurons in (I) KO-*Cdk5r1*; MADM-4 (*MADM-4^GT/TG^;Cdk5^-/-^;Emx1^Cre/+^*), (J) *Dab1*-MADM-4;KO-*Cdk5r1* (*MADM-4^GT/TG,Dab1^;Cdk5r1^-/-^;Emx1^Cre/+^*), and (G) double-KO *Dab1*-MADM-4;KO-*Cdk5r1* (*MADM-4^GT,Dab1/TG,Dab1^;Cdk5r1^-/-^;Emx1^Cre/+^*). Relative distribution (%) of MADM-labeled projection neurons is plotted in ten equal bins across the cortical wall. (H) Direct distribution comparison of *Dab1^-/-^* mutant cells upon sparse (grey) and global (black) KO in *Cdk5r1^-/-^* mutant background. All analysis was carried out at P21. n=3 for each genotype. From each animal 10 (MADM-11) or 20 (MADM-4) hemispheres were analyzed. Nuclei were stained using DAPI (blue). Data indicate mean ± SD, *p<0.05, **p<0.01, and ***p<0.001. Scale bar: 100μm.

### Tissue-wide non-cell-autonomous effects impacting on neuronal migration are specific for distinct signaling pathways

Global, but not sparse KO, of *Dab1* results in tissue-wide effects affecting the single cell *Dab1^-/-^* mutant phenotype in quite a similar manner to the above p35/CDK5 findings. Whether the observed tissue-wide effects exhibit ‘gene’ specificity or may dominate over distinct signaling pathways was however not clear. We thus conceived epistasis experiments (Figure S6) to test possible pathway exclusivity of tissue-wide properties upon global gene KO. First we utilized MADM-11 platform to generate Control, global and sparse *Cdk5r1* deletion paradigms in a full *Dab1^-/-^* KO background (Figures 6F-6H). We thus tested whether tissue-wide effects due to global *Dab1* ablation may affect or interfere with the cell-autonomous *Cdk5r1^-/-^* mutant phenotype (i.e. accumulation of mutant cells in lower layers and the white matter as described in Figure 1F). We however observed a similar cell-autonomous *Cdk5r1^-/-^* mutant phenotype (upon sparse mosaic *Cdk5r1* deletion) in *Dab1^-/-^* background (Figure 6F) as in *Dab1^+/+^* background (i.e. in *Cdk5r1*-MADM context). Next we reversed the genetic background conditions and utilized MADM-4 platform to generate Control, global and sparse *Dab1* deletion paradigms in a full *Cdk5r1^-/-^* KO background (Figures 6I-6L). Again, the cell-autonomous *Dab1^-/-^* mutant phenotype upon sparse deletion was highly similar in *Cdk5r1^-/-^* KO (Figure 6J) when compared to *Dab1*-MADM in *Cdk5r1^+/+^* background (Figure 6B). In contrast to the above findings, concomitant global KO of both *Cdk5r1* and *Dab1* resulted in a relatively uniform distribution of double mutant cells across the cortical wall (Figures 6G and 6K), comparable to the distribution in individual KO of *Dab1* (Figure 6E) or *Cdk5r1* (Figure 6I), respectively. In summary, tissue-wide non-cell-autonomous effects predominate the cell-autonomous phenotype albeit in a gene-specific manner.

### Global KO of both *Cdk5r1* and *Dab1* triggers deregulation of genes associated with ECM and cell adhesion

The phenotypic manifestation of global tissue-wide effects appeared very similar for *Cdk5r1* and *Dab1* because global ablation of either gene resulted in relatively uniform distribution of mutant neurons across the vertical axis of the cortical wall. We therefore asked next whether we might find overlap in deregulated gene expression upon global *Cdk5r1* and *Dab1* LOF or not. To this end we applied the identical approach as described above and FACS-isolated green GFP^+^ MADM-labeled cells (using MADM-11 platform) from control-MADM (*MADM-11^GT/TG^;Emx1^Cre/+^*) and MADM;KO-*Dab1* (*MADM-11^GT/TG^;Dab1^-/-^;Emx1^Cre/+^*) at P0 (Figure 7A). .We complemented this dataset with control-MADM (*MADM-11^GT/TG^;Emx1^Cre/+^*) and KO-*Cdk5r1*-MADM (*MADM-11^GT,Cdk5r1/TG,Cdk5r1^;Emx1^Cre/+^*) data as described before (Figure 5). We performed DEG analysis on the combined data set (see also Materials and Methods). *Dab1* expression level was close to zero in MADM;KO-*Dab1* but readily detectable in all other samples, validating the MADM;KO*-Dab1* ablation paradigm (Figure S7A).

**Figure 7.**
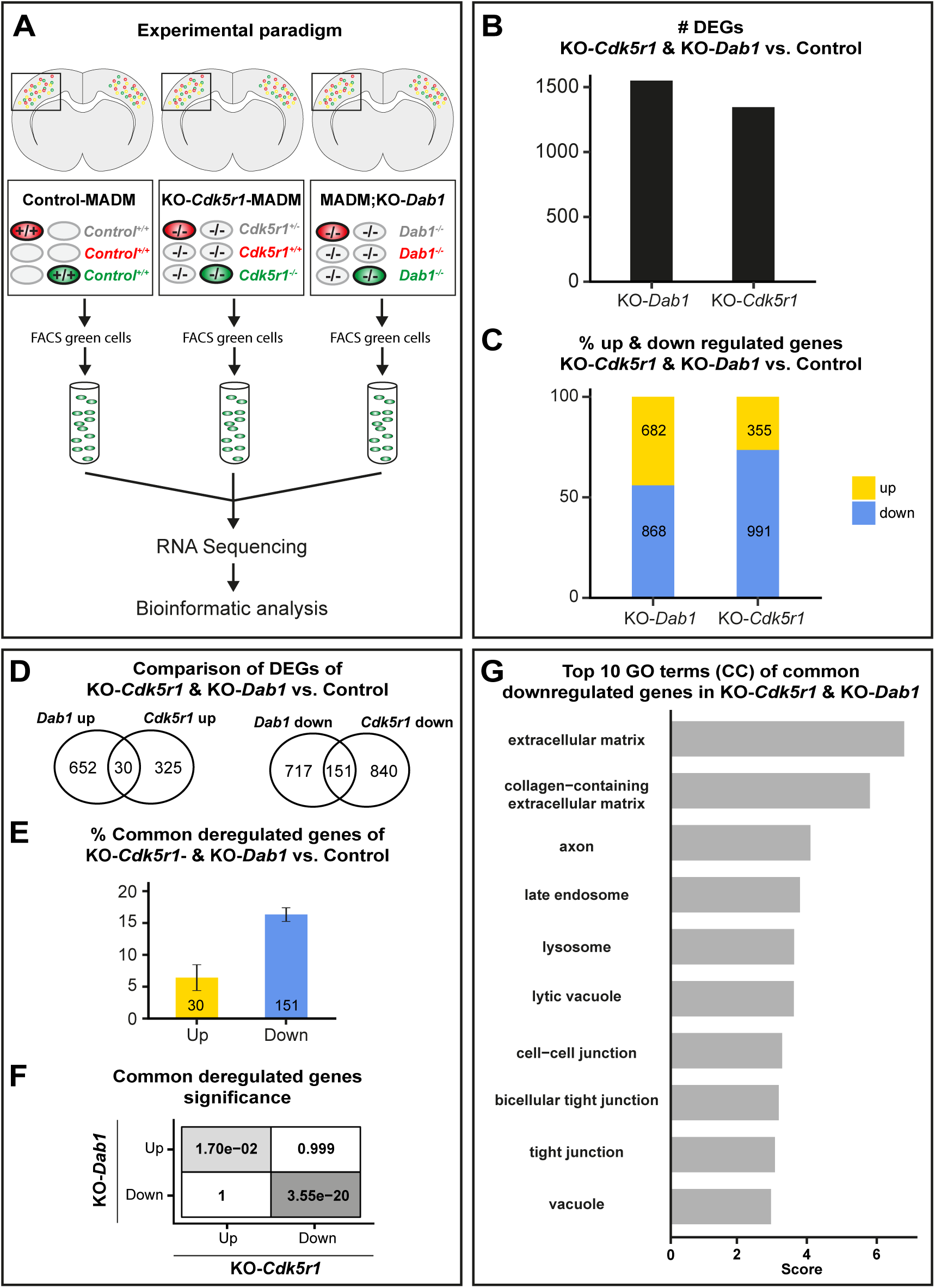
Gene expression upon combined global KO of *Cdk5r1* and *Dab1*. **(A)** Experimental paradigm and pipelines for gene expression profiling in Control-MADM (left), KO-*Cdk5r1*-MADM (middle; KO-*Cdk5r1*), and MADM;KO-*Dab1* (right; KO-*Dab1*) at P0. **(B)** Number of differentially expressed genes (DEGs) in KO-*Cdk5r1* and KO-*Dab1* versus Control. **(C)** Percentage of up- and downregulated genes in KO-*Cdk5r1* and KO-*Dab1* versus Control. **(D)** Venn diagrams showing common up- and down-regulated genes in KO-*Cdk5r1* and KO-*Dab1* versus Control. **(E)** Percentage of common up- and downregulated genes in KO-*Cdk5r1* and KO-*Dab1* versus Control. **(F)** Significance of all pairwise overlaps of DEGs shown in (D). **(G)** Top 10 GO terms of commonly downregulated genes in KO-*Cdkr5r1* and KO-*Dab1*, according to overlap shown in (D, right). Commonly upregulated genes did not yield any significant GO term enrichment.

Next, we analyzed DEGs in KO-*Cdk5r1-*MADM and MADM;KO-*Dab1*, relative to Control, and compared the data to each other. Upon global ablation of either, *Cdk5r1* or *Dab1*, we found more than 1000 deregulated genes in both mutants (Figure 7B) with the majority of DEGs being downregulated (Figure 7C). We next analyzed the overlap of DEGs in *Cdk5r1* and *Dab1* KOs and found both common and non-common deregulated genes (Figure 7D). The majority of the commonly deregulated genes were downregulated (Figure 7E) and the overlap of both, up- and downregulation was significant (Figure 7F). Among the upregulated genes we did not obtain any significantly enriched GO-terms although the non-common deregulated genes showed overlap of many of their respective GO-terms between KO-*Cdk5r1-* MADM and MADM;KO-*Dab1* (Figure S7B). In contrast, among the commonly downregulated genes, we found significant GO-terms associated with cell-cell and cell-matrix interaction (Figure 7G). Taken together, the above analysis suggested that cell-adhesion and ECM represent major elements in tissue-wide effects upon global KO of genes encoding components of the p35/CDK5 and Reelin/DAB1 signaling pathways.

## DISCUSSION

Radial migration of cortical projection neurons has been extensively studied and a rich catalogue of regulatory signaling cues and pathways have been compiled. The systematic study of the cell-autonomous functions of the genes encoding the signaling cascades, have revealed an extensive genetic framework. However, the nature and relative contributions of global tissue-wide effects, in regulating radial migration in the developing neocortex, remained unclear. To this end we established genetic paradigms enabling the dissection of cell-autonomous gene function and non-cell-autonomous effects. Our data revealed that the genetic landscape of the cellular environment, surrounding migrating neurons, fundamentally impacts the individual single cell phenotype (Figure 8).

**Figure 8.**
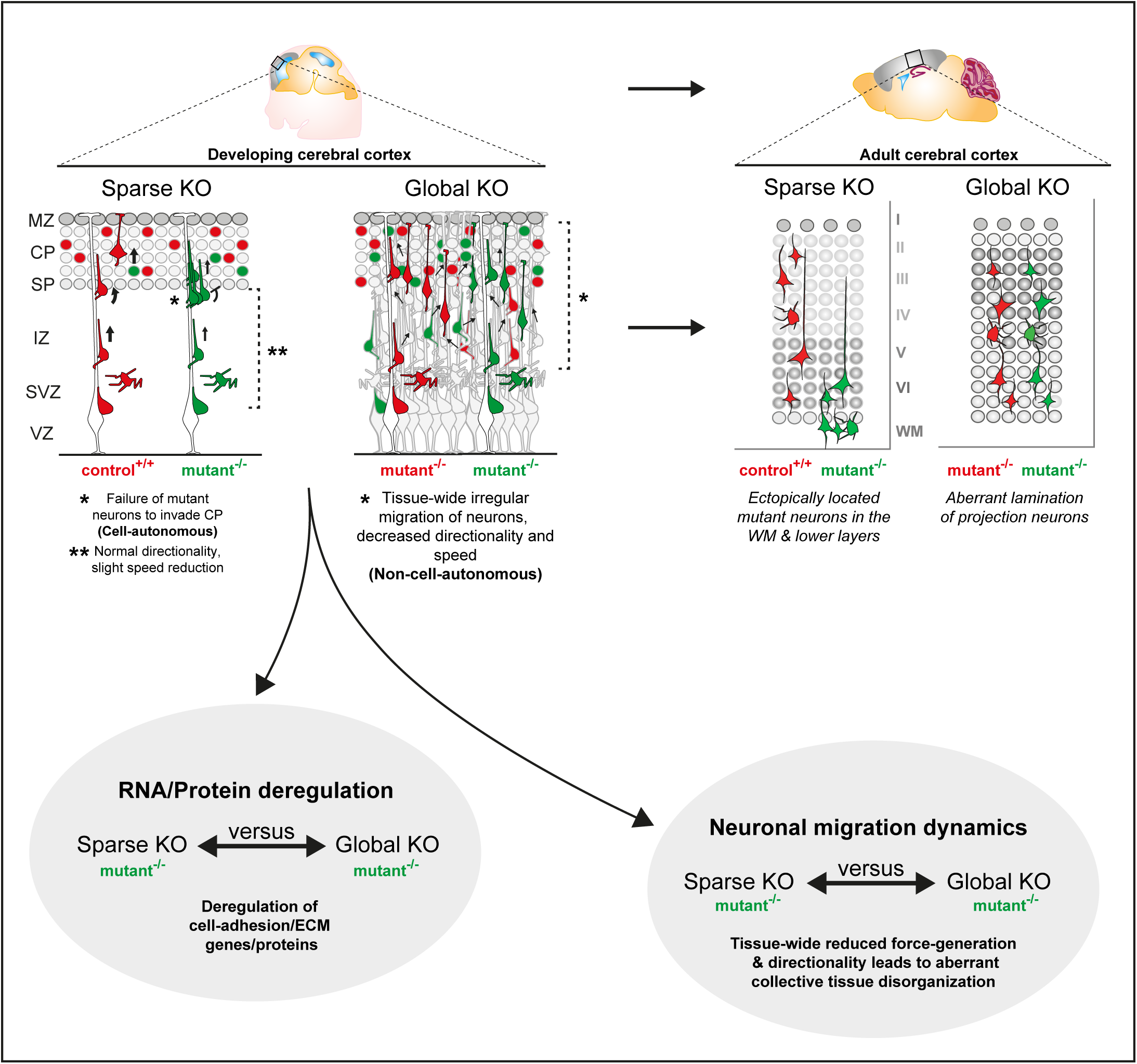
Interplay of cell-autonomous and global tissue-wide properties in cortical projection neuron migration. Schematic illustrating the MADM-based subtractive phenotypic analysis of sparse genetic mosaics (control background) and global knockout (cKO/KO) (mutant background), both coupled with fluorescent MADM-labeling of homozygous mutant and control neurons. Such assay enabled the high-resolution analysis of projection neuron migration dynamics in distinct genetic environments with concomitant isolation of genomic and proteomic profiles. In combination with computational modeling, we utilized these experimental paradigms to visualize non-cell-autonomous effects in radial neuron migration at single cell resolution. In sparse KO, mutant neurons migrated more dynamically and expressed cell adhesion molecules similar like in Control. However, in global KO, we observed that cell adhesion molecules were significantly downregulated. Mutant neurons in global KO also showed much more severe migration phenotype resulting in drastic disorganization of mature cortical wall.

We focused our analysis mainly on the well characterized p35/CDK5 pathway providing a defined genetic framework and context. We could show that sparse p35/CDK5 ablation results in a highly specific phenotype that we attributed to the loss of cell-autonomous gene function. Individual *p35/Cdk5^-/-^* mutant cortical projection neurons in sparse genetic mosaic conditions failed to pass the border between the IZ and the emerging CP and progressively accumulated below the CP and the white matter. However, radial migration from the VZ and through the IZ occurred, albeit at lower speed. In contrast, single *p35/Cdk5^-/-^* mutant neurons in a global homozygous *p35/Cdk5^-/-^* mutant KO environment migrated even slower in the VZ/IZ, and showed skewed directionality. The perhaps more surprising finding was that in the above full KO condition, *p35/Cdk5^-/-^* mutant neurons distributed relatively evenly across the entire cortical wall. Thus the cell-autonomous phenotype as observed in sparse KO scenario seemed wiped-out. There are a few (not mutually exclusive) possibilities that could explain the distinct phenotypic observations. First, the IZ-CP border lost its property as a physical gatekeeper upon global loss of *p35/Cdk5*. Such scenario actually formed an important basis of our modeling approach where we assumed uniform tissue stiffness/resistance that radially migrating neurons encounter upon global p35 ablation. Indeed, the simulation of such conditions revealed very similar migration trajectories of *p35/Cdk5^-/-^* mutant cells as observed by live-imaging in experimental global *p35/Cdk5* KO conditions. Second, *p35/Cdk5^-/-^* mutant neurons in sparse genetic mosaic lost an essential intrinsic property required for transiting the IZ-CP border, reflecting the cell-autonomous *p35/Cdk5* gene functions. One important p35/CDK5 downstream signaling hub is NDEL1 which is phosphorylated at specific sites by CDK5 (Niethammer et al., 2000; Sasaki et al., 2000). Interestingly, sparse mosaic deletion of *Ndel1* results in a congeneric phenotype, to the sparse *p35/Cdk5^-/-^* condition, whereby *Ndel1^-/-^* mutant neurons fail to invade the CP and progressively accumulate below the CP and later in the white matter (Hippenmeyer et al., 2010). However, tissue-wide *Ndel1* KO results in severely disorganized neocortex with seemingly immobile *Ndel1^-/-^* mutant neurons (Youn et al., 2009). Thus, sparse loss of any component of the p35/CDK5/NDEL1 signaling hub leads to the inability of individual radially-migrating neurons to enter the target area, the emerging CP, while global KO results in predominant cortical tissue disorganization with slower or even immobile projection neurons. The sparse ablation of p35/CDK5/NDEL1 might lead to deficits in LIS1/Dynein-mediated force generation (Bradshaw and Hayashi, 2017; Jheng et al., 2018; Marín et al., 2010; Reiner and Sapir, 2013; Vallee et al., 2009) necessary to overcome the increased physical tissue resistance at the IZ/CP border, although the precise biochemical mechanism remains to be clarified in future studies.

The observation that global KO, of the same signaling component as in sparse KO, results in predominant tissue-wide effects appears to exhibit pathway specificity. As such, sparse elimination of *p35* gene function in a *Dab1^-/-^* global KO background (which also shows severe disorganization of the cortical wall) showed a similar cell-autonomous phenotype (failure to enter CP) like in *Dab1^+/+^* background conditions. Thus, the cell-autonomous *p35^-/-^* phenotype was preserved regardless of the global *Dab1* genotype. It will be revealing to investigate the generality of the above findings in future experiments to systematically assess potential epistatic interactions of mutations in distinct genes that lead to cortical malformation.

What are the underlying characteristics of global tissue-wide effects impacting on radial projection neuron migration? It is clear that any tissue consists of a complex extracellular environment. Thus individual cells are exposed to many extrinsic elements including 1) secreted factors acting locally, globally or even systemically; 2) the extracellular matrix; and 3) neighboring cells mediating cell-cell interactions through receptors and/or direct physical stimuli (Hansen and Hippenmeyer, 2020). While secreted factors mostly signal to other cells (besides autocrine signaling), and thus mainly act non-cell-autonomously, how cell-intrinsic signaling molecules could act at the global tissue-wide level is not yet clear. Our MADM data is based on the genetic deletion of the intracellular CDK5 kinase or DAB1 adapter protein function. In both instances the cell-autonomous migration phenotype upon sparse deletion differed from the one upon global KO. In order to get insights at the molecular level we pursued transcriptome/proteome analysis and found that upon sparse deletion of for instance p35, very few genes and proteins were deregulated in mutant cells. In stark contrast, global KO let to a high number of deregulated proteins and genes. A sizeable fraction of differentially expressed genes even correlated significantly with deregulation of their encoded proteins. The top GO terms associated with deregulated genes/proteins included ECM, receptors and cell adhesion. These results indicate that at the global tissue-wide level the ECM and cell adhesion landscape was drastically changed in global KO condition when compared to wild-type. Although p35/CDK5 and DAB1 signal in parallel, deregulated ECM and cell adhesion was a common denominator upon global KO. Our data are in agreement with earlier studies that showed N-cadherin dependent signaling to be relevant for both p35/CDK5 (Kwon et al., 2000) and DAB1 (Franco et al., 2011; Sekine et al., 2011, 2012) pathways, respectively. Interestingly, transcriptomic analysis in whole cortex KO mouse models for *Ndel1*, *Lis1*, and *Ywhae*, all acting in the LIS1/Dynein signaling pathway and to some extent downstream of p35/CDK5, have also revealed altered cell adhesion and cytoskeleton organization pathways (Pramparo et al., 2011).

Migrating neurons can exert positive and negative interaction on each other depending on the cellular environment and their genetic constitution (Gorelik et al., 2017; Greenman et al., 2015; Hansen and Hippenmeyer, 2020). In collective cell migration for instance, cell-cell interactions balancing adhesion and repulsion is a key mechanism (Shellard and Mayor, 2020). Besides, direct physical interactions might be critical whereby for example less agile mutant cells get passively pushed, pulled, or simply piggyback on more dynamic cells.

Our findings put a new perspective on the clinical symptoms and/or appearance of disease-causing mutations affecting radial migration in particular, but also mutations causing cortical malformation in general. For instance in focal malformations of cortical development (FMCD) a small fraction of mutant cells can disrupt neighboring cells and even large areas thereby affecting overall cortical architecture (Baek et al., 2015; Lee et al., 2012; Poduri et al., 2012; Rivière et al., 2012). In order to obtain a better understanding of FMCD disease etiology it will thus be essential to rigorously scrutinize the contribution of not only cell-autonomous loss of gene function but also tissue-wide and systemic effectors. Interestingly, lissencephaly with cerebellar hypoplasia (LCH) has been attributed to individuals with mutations in *CDK5* or *RELN* (Hong et al., 2000; Magen et al., 2015). The common clinical appearance in such patients could (at least in part) emerge from tissue-wide effects alike the ones we observed in our MADM-based analysis of *Cdk5r1/CDK5* and *Dab1*, respectively.

## CONCLUSION

Our study provides quantitative evidence that global tissue-wide effects play essential roles in the control of radial projection neuron migration in the developing cortex. Based on defined genetic conditions in combination with single cell tracing we could show that sparse mosaic ablation of gene function results in highly specific migration phenotypes. In contrast, in global KO, individual migrating neurons exhibit distinct deficits that result from predominant tissue-wide effects. Altogether, cortical projection neuron migration is tightly regulated by intrinsic gene function and depending on the cellular and genetic landscape of the overall surrounding tissue.

## ACKNOWLEDGMENTS

We thank A. Sommer and C. Czepe (VBCF GmbH, NGS Unit), L. Andersen, J. Sonntag, and J. Renno for technical support and/or initial experiments; M. Sixt, J. Nimpf and all members of the Hippenmeyer lab for discussion. This research was supported by the Scientific Service Units (SSU) of IST Austria through resources provided by the Imaging & Optics Facility (IOF), Lab Support Facility (LSF), and Preclinical Facility (PCF).

## STUDY FUNDING

A.H.H. was a recipient of a DOC Fellowship (24812) of the Austrian Academy of Sciences. This work also received support from IST Austria institutional funds; the People Programme (Marie Curie Actions) of the European Union’s Seventh Framework Programme (FP7/2007-2013) under REA grant agreement No 618444 to S.H.

## APC FUNDING

APC funding was obtained by IST Austria institutional funds.

## AUTHORS’ CONTRIBUTIONS

S.H. and A.H.H. conceived the research. A.H.H., C.St., and A.H generated all experimental MADM tissue. A.H.H., C.St., T.R, and S.L., performed all the experiments except LCMS which was performed by A.N. A.H.H., F.M.P, and C.So. analyzed imaging data. F.M.P. performed computational and bioinformatics analysis of RNA-seq and proteomics with inputs from A.H.H. and S.H. M.R. established the computational model with input from B.H, A.H.H and S.H. L.H.T provided critical resources. S.H. and A.H.H. wrote the manuscript with input from F.M.P. and M.R. All authors edited and proofread the manuscript.

## MATERIALS AND METHODS

### Mouse Lines

All mouse colonies were maintained in accordance with protocols approved by the institutional animal care and use committee, institutional ethics committee, and the preclinical core facility (PCF) at IST Austria. Experiments were performed under a license approved by the Austrian Federal Ministry of Science and Research following the Austrian and EU animal laws (license numbers: BMWF-66.018/0007-II/3b/2012 and BMWFW-66.018/0006-WF/V/3b/2017).

Mice with specific pathogen-free status according to FELASA recommendations (Mähler (Convenor) et al., 2014) were bred and maintained in experimental rodent facilities (room temperature 21 ± 1°C [mean ± SEM]; relative humidity 40%–55%; photoperiod 12L:12D). Food (V1126, Ssniff Spezialitäten GmbH, Soest, Germany) and tap water were available ad libitum.

Mouse lines with MADM cassettes inserted on chr.4, chr.5, and chr.11 (Contreras et al., 2021; Hippenmeyer et al., 2010) [MADM-4-GT, MADM-4-TG, MADM-5-GT, MADM-5-TG, MADM-11-GT (JAX stock # 013749), MADM-11-TG (JAX stock # 013751)], *Cdk5r1* (Chae et al., 1997) (JAX stock # 004163), *Cdk*5-flox (Samuels et al., 2007) (JAX stock # 014156), *Dab1* (Howell et al., 1997) (JAX stock # 003581); *Emx1*-Cre (Gorski et al., 2002) (JAX stock # 005628) were previously described. We have not observed any influence of sex on the results in our study, and all experiments and analyses were carried out using animals of both sexes. Phenotypic time course analysis of *Dab1*-MADM-4, *Cdk5*-MADM-5, *Cdk5r1*-MADM-11 in combination with *Emx1*-Cre was performed at E14, E16, P0, and P21. For sequencing and/or proteomics experiments, MADM-11 animals were used in combination with *Emx1*-Cre and were analyzed at E13, E16, and P0. Genetic epistasis experiments of *Cdk5r1* and *Dab1* on MADM-4 and MADM-11 in combination with *Emx1*-Cre were all performed at P21.

### Isolation of fixed tissue

Tissues from postnatal time points (P15/P21) were collected by cardiac perfusion. Mice were deeply anesthetized through injection of a ketamine/xylazine/acepromazine solution (65 mg, 13 mg, and 2 mg/kg body weight, respectively) and unresponsiveness was confirmed through pinching the paw. The diaphragm of the mouse was opened from the abdominal side to expose the heart. Cardiac perfusion was performed with PBS followed immediately by ice-cold 4% PFA prepared in PB buffer (Sigma-Aldrich). Brains were removed and further fixed in 4% PFA for 24 hours at 4°C to ensure complete fixation. Brains were cryopreserved with 30% sucrose (Sigma-Aldrich) solution in PBS for approximately 48 hours. Brains were then embedded in Tissue-Tek O.C.T. (Sakura). For adult time points, 45μm coronal sections were collected in 24 multi-well dishes (Greiner Bio-one) and stored at −20°C in antifreeze solution (30% v/v ethylene glycol, 30% v/v glycerol, 10% v/v 0.244M PO4 buffer) until used. Tissue from embryonic time points (E14/ E16) and postnatal day zero (P0) was directly transferred into 4% PFA and kept at least 24 hours at 4°C. Cryopreservation and embedding were done as described for adult brains. For embryonic and early postnatal brains 25μm cryosections were directly mounted onto Superfrost glass-slides (Thermo Fisher Scientific).

### Immunohistochemistry

Brain sections were mounted onto Superfrost glass-slides (Thermo Fisher Scientific) and let to dry, followed by 3 wash steps each of 5 minutes with PBS. Tissue sections were blocked for 30 minutes in a buffer solution containing 5% normal donkey serum (Thermo Fisher Scientific) and 0.5% Triton X-100 in PBS. Primary antibodies for GFP (Chick, Aves Labs Inc.) and RFP (Rabbit, MBL) were mixed in the blocking buffer and incubated on the tissue for at least 12 hours at 4°C. Sections were washed 3 times for 5 minutes each with PBT (0.5% Triton X-100 in PBS) and incubated with the corresponding secondary antibodies Alexa Fluor 488 (Anti-Chicken IgG, Jackson ImmunoResearch Labs) and Cy3 (Anti-Rabbit IgG, Jackson ImmunoResearch Labs) diluted in PBT for 1 hour. Sections were then washed 2 times with PBT and once with PBS each for 5 minutes. Finally, nuclear staining was done using 10 minutes of incubation with PBS containing 2.5% DAPI (Thermo Fisher Scientific). Sections were embedded in mounting medium containing 1,4-diazabicyclooctane (DABCO; Roth) and Mowiol 4-88 (Roth) and stored at 4°C.

### Imaging of fixed brain tissue

Histological brain sections were imaged using confocal microscopy (Zeiss inverted LSM800) or epifluorescence microscopy (Olympus VS120 Slide scanner). Confocal images were recorded on a Zeiss LSM 800 laser-scanning confocal microscope mounted with a plan-apochromat 10x/0.45, WD=2.1 mm objective. Excitation/emission wavelengths were 488/509 nm (EGFP), 554/581nm (tdTomato), and 353/465nm (DAPI). Z-series images were collected on a PC running ZEN 2.6 software (Zeiss). Image series were Z-projected, stitched and contrast-enhanced using ZEN 2.6 software (Zeiss). Slidescanner images were recorded with a 10x / NA 0.4 objective. Excitation/emission wavelengths were 485/518 nm (FITCH), 560/580nm (Cy3), and 387/455nm (DAPI). Slide scanner images were processed using ImageJ software (Schindelin et al., 2012).

### Analysis of relative distribution of MADM-Labeled neurons

Images were imported into ImageJ software (Schindelin et al., 2012) and MADM-labeled neurons were manually quantified based on the respective fluorescent marker expression and their relative position, which was calculated with respect to the bottom of the ventricle and the pial surface (for details see http://github.com/hippenmeyerlab/cell2layer). The analysis script used, computes the relative and absolute distances of each manually marked neuron to its boundaries (ventricular surface and pia) provided manually as two segmented lines. For each neuron the shortest distance to the two-layer boundaries is computed, resulting in two distances d_1_ and d_2_. The normalized (relative) distance is computed by:

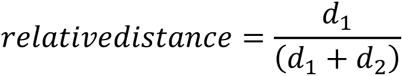

Statistical analyses were done with GraphPad Prism 8.0.1, applying an arcsin conversion of relative percentages, a two-way ANOVA, and a Tukey post hoc test.

### Slice culture and time-lapse imaging

Embryos were collected at E16 and stored in ice-cold PBS during genotyping. Immediately after genotyping, MADM labeled embryonic brains were dissected and mounted in 4% low-melting agarose (Fisher BioReagents). 300µm coronal slices were prepared in oxygenated ice-cold artificial cerebrospinal fluid (ACSF) using a vibratome (Leica VT 1200S). Thereafter, slices were placed on Milicell culture inserts (Millipore) in 6-well glass-bottom dishes (MatTek) containing culture medium (1% 100X N2 supplement (Gibco), 1% Penicillin-Streptomycin (Gibco) in transparent F12/DMEM (Gibco)) and incubated (37°C, 5% CO2) for at least 45min prior to imaging acquisition. To reduce the evaporation of media during imaging a FoilCover lid (Pecon) was applied on top of the glass-bottom dishes during time-lapse imaging. A time-lapse of minimum 15hrs with a framerate of 15± min was recorded unidirectionally at 7 Z-positions with 5μm spacing using confocal microscopy [(Zeiss LSM800, Plan-Apochromat 10x/0.45, WD=2.1 mm objective, equipped with a heating chamber and stage-top incubator chamber & gas mixed (Ibidi) (37°C, 5% CO2)]. Excitation/emission wavelengths were 488/509 nm (EGFP) and 554/581nm (tdTomato). Time-lapse images were collected on a PC running Zeiss ZEN Blue software. Time-lapse image series were Z-projected, time-stitched using ZEN blue software.

### Correction of non-linear local drift in time-lapse images

To correct any local tissue drift in the original 3D multi-channel movies, we developed the Python package undrift. First, dense optical flow from successive image pairs is estimated with the Farnebäck method (Farnebäck, 2003) using the OpenCV library (version 3.3.1). For movies with more than one input channel, we used the averaged channel intensities before estimating the optical flow for each pixel. Input parameters for the Farneback method were set as follows: the number of image pyramid levels to 3, the averaging window size to 512x512 (px), the size of the pixel neighborhood used to find polynomial expansions to 5 px, the standard deviation of the Gaussian that is used to smooth derivatives to 0.4 px and the number of iterations per pyramid level to 3. Parameters were optimized to capture the movement of single cells and the locally coherent drift of tissue regions (if present). Then, the pairwise optical flow fields were smoothed (locally weighted averaged) with a spatio-temporal Gaussian (σ_t=1 px and σ_xy=25.6 px) using the scikit-image library (0.16.2). The strong spatial smoothing effectively removes movement on a small scale (single cells), whereas spatially coherent optical flow on a bigger scale (tissue drift) is maintained in the output. The smoothed pairwise optical flow fields are integrated over time to obtain an optical flow field relative to the reference frame (first time-point) using cubic spline interpolation and the Python SciPy library (version 1.4.1). New movies are rendered by artificially unwarping this integrated flow field on the original movie channels starting from the reference frame. For more information see http://github.com/hippenmeyerlab/undrift.

### Analysis of Neuronal Trajectories

Neurons were tracked semi-automatically with the ImageJ plugin TrackMate (Tinevez et al., 2017) using the LoG detector (estimated blob diameter: 10.0micron, threshold: 2.0, Median filter: enabled, sub-pixel localization: enabled) and the linear motion LAP tracker (initial search radius: 15, max search radius: 15, max frame gap: 2) for each channel. Tracks were manually curated to ensure correct tracking of neurons. Only red and green neuronal tracks were included in the analysis, all yellow neurons were excluded in the analysis. Tissue compartments (upper/lower bin) were drawn manually. All parameters were extracted in a .csv file for analysis. For each neuron, we first determined if it was located in the located in the upper / lower bin in the first frame or last frame. In Figures 3E-3F: Each cell-track was grouped into Upper Bin if in any frame the cell was positioned in the Upper Bin area. The cell-track was grouped into Lower Bin if the cell was in no frame placed in the Upper Bin area. An arcsin conversion were performed of relative percentages for statistical calculations. Figures 3G-3H: We extracted the x/y coordinates of cells based on their position (Upper Bin, Lower Bin). The resulting cells were re-grouped into cell-tracks and for each track, we calculated mean straight-line speed and directionality as follows.

Distance of one cell between two frames as:

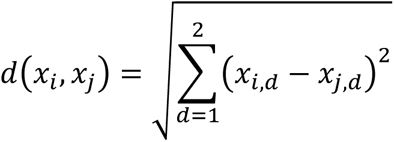

Sum of all distances (total distance traveled) with N being the number of frames a cell was tracked in:

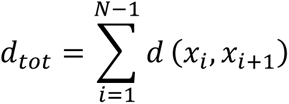

Net distance traveled:

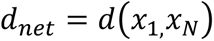

Net time traveled:

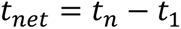

Mean straight line speed:

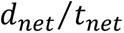

Directionality (meandering) index:

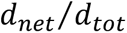

Note that the time between frames can differ for each cell (for example if a cell was not identified in one frame), and that the total time a cell was tracked can also differ between cells. Therefore, to assure the same framerate for the analysis, we calculated the velocities in micrometer per minute. Note that a single track can be divided into sub-tracks if crossing the middle line. These tracks are ‘crossing’ and make up on average ∼6% of all tracks. For 3E-3H we ranked each cell-track from each video based on d_net_ and plotted the indicated values for the top 15 cells for each video. Statistical analyses were done with GraphPad Prism 8.0.1, applying a two-way ANOVA and a Tukey post hoc test.

### Formulation of computational migration model

The formulated 2D migration model is evaluated via a Python 3.6 script. The force generation mechanism is based on a random walk model with directional bias on a single cell of unit mass (Caffrey et al., 2014). We include an alteration from Caffrey et al., 2014 by generating a 2D force *F*_*gen*_ in an intermediate step instead of directly calculating a displacement for each cell. The total force acting on a given cell at each time instance is given by:

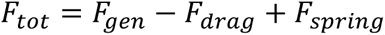

For each timestep *dt* the internal molecular machinery of a cell generated force *F*_*gen*_. If this force is insufficiently high to overcome the subsequently resulting drag force *F_drag_* it is stored in form of a spring-like force *F_spring_* for the next timestep. Whereas the drag force is defined as:

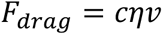

Here, *η* denotes the dynamic viscosity and c is a parameter dependent on the cell shape, which we consider a spherical particle of unit radius. In our model both parameters are considered to be locally constant and can therefore be unified as a resistance parameter *R*. Similar models have been previously described for 3D cell migration in extra cellular matrices (Zaman et al., 2005). We include the cell intrinsic difference between Control and KO-*Cdk5r1*-MADM in the form of a directionality parameter *ρ* and a force scaling parameter *α* (Caffrey et al., 2014). Here, a value of *ρ* = 0.5 corresponds to a pure random walk model, which KO-*Cdk5r1*-MADM neurons tend towards. However, Control neurons are closer to *ρ* = 1, indicating a high directionality. The force scaling parameter *α* controls the magnitude of the generated force and henceforth velocities. The non-cell-autonomous effect is included by introducing a linear coupling of directionality and force generation coefficient with the ratio *β* of control cells (*N_Control_*) to mutant cells (*N*_*KO*_ ). Boundary values for *α* and *ρ*at *β* = 0 and *β* = 1 correspond to values from experimental data for KO-*Cdk5r1*-MADM and Control, respectively. For more details of our model and parameters see Table 1 below and Figure S3.

**Table 1.** Modelling Parameters.

|  |  |
| --- | --- |
| Directionality: | Force scaling: |
| $\rho = \begin{cases} 0.51 & \beta < 0.1 \\ f(\beta) & 0.1 < \beta < 0.05 \\ 0.65 & \beta > 0.05 \end{cases}$ | $\alpha = \begin{cases} 0.53f_{Ctrl} & \beta < 0.1 \\ f(\beta)f_{Ctrl} & 0.1 < \beta < 0.05 \\ 0.9f_{Ctrl} & \beta > 0.05 \end{cases}$ |
| Resistance Control environment | Resistance mutant environment |
| $R_{Ctrl,tissue} = \begin{cases} R_1 = 5 \cdot 10^3 & y < 0.45 \\ R_2 = 12 \cdot 10^5 & y \leq 0.65 \\ R_3 = 4 \cdot 10^3 & y > 0.65 \end{cases}$ | $R_{KO,tissue} = 7 \cdot 10^3$ |

### Preparation of Cell Suspension and FACS

Preparation of cell suspension for cell sorting was prepared as previously described (Laukoter et al., 2020b) for E13, E16, and P0 time points for RNA-seq and P0 for proteomics. From the overall MADM samples, GFP^+^ cells were collected for each genotype. For RNA-sequencing, cells were sorted directly into a custom-made lysis buffer (30nM TRIS pH8, 10nM EDTA pH8, 1% SDS and 200μg/μL Proteinase K). For proteomics, cells were prepared as described above except no serum was added to the media and an extra wash step with DMD/F12 wash was carried out. Samples for proteomics were sorted directly into 50ul lysis buffer (LYSE-NHS, Preomics iST-NHS kit).

### RNA Extraction of MADM Samples for RNA Sequencing

Directly after cell sorting using FACS, samples were incubated for 30min at 37°C. Total volume was filled to 250μl using RNase-free H2O (Thermo Fisher Scientific) followed by the addition of 750μl Trizol LS (Thermo Fisher Scientific). Samples were mixed by inversion (5 times). After a 5min incubation step at RT, the entire solution was transferred into a MaXtract tube (QIAGEN). 200μl chloroform (Sigma-Aldrich) was added, followed by 3 times 5sec vortexing and 2min incubation at RT. Samples were centrifuged for 2min at 12000rpm at 18°C. The supernatant was transferred to a new tube and isopropanol (Sigma-Aldrich) was added in a 1:1 ratio. For better visibility of the RNA pellet 1μl GlycoBlue (Thermo Fisher Scientific) was added and the entire solution was mixed by vortexing (3x 5sec). Samples were left for precipitation o/n at −20°C. After precipitation samples were centrifuged for 20min with 14000rpm at 4°C. The supernatant was removed and the RNA pellet was washed with 70% ethanol, followed by a 5min centrifugation step (14000rpm at 4°C). The RNA pellet was resuspended in 12,5μl RNase-free H_2_O. RNA quality was analyzed using RNA 6000 Pico kit (Agilent) following the manufacturer’s instructions. The RNA samples were stored at −80°C until further use. RNA sequencing was performed by VBCF GmbH on Illumina platforms.

### Statistical Analysis of RNA-Seq

Read processing, alignment and annotations are described elsewhere (Laukoter et al., 2020b). STAR alignment parameters: clip5pNbases 3, outFilterMultimapNmax 1, --outSAMtype BAM SortedByCoordinate and quantMode GeneCounts. Downstream analyses were performed in R (v3.6.1). Read coverages of the deleted *Cdk5r1* region (chr11:80477417-80478722, mm10) and the deleted *Dab1* region (chr4:104605298-104605437, mm10) were calculated using bedtools intersect with the -split option on the aligned bam file produced by STAR. These read counts were added to the count tables produced by STAR with the gene name *Cdk5r1*_del and *Dab1*_del respectively.

For Figures 5A-5E we analyzed 81 samples and removed 9 samples with a low percentage of uniquely aligned reads (<50%) or due to their position on the PCA plot. Statistics on differential expression between all pairs of genotypes were calculated with DESeq2 (v1.26.0) using contrasts for each developmental time point separately. To reduce noise only genes with an average read coverage of >10 were used in the analyses. We used an adjusted p-value (padj) cutoff of 0.05 for differential expressed genes (DEG) for all analyses in Figures 5A-5E Up- and down-regulated genes were determined by a log_2_ fold-change of >0 or <0 respectively. For Gene Ontology term enrichment we used the enrichGO function from the clusterProfiler package (v3.14.0) with parameters: universe = [all informative genes in the respective comparison], ont = "ALL", pool = T, readable = T, OrgDb = org.Mm.eg.db (v3.10.0), minGSSize = 20, maxGSSize = 500, pvalueCutoff = 0.1, qvalueCutoff = 0.2. For the GO term plot in Figure 5E we focused on P0 timepoint and first removed GO terms that are not related to neuronal development by removing GO terms with blood, vascu, or angio in their description. Then we calculated the negative log10 of the uncorrected p-value for the remaining GO terms (score). Finally, we ranked GO terms by the score and plotted the score of the top 10 GO terms focusing on the ontology BP.

For Fig. 7 we analyzed 38 samples consisting of 22 samples already used for Figures 5A-5E (control, *KO-Cdk5r1-MADM*, P0 time point) and 16 new samples (control, *Dab1*-KO). We removed 3 samples due to their position on a PCA plot. Statistics on differential expression were calculated as for Fig. 5A-5E using genes with an average read coverage over all samples > 20. For all analyses in Fig. 7 we used an adjusted p-value (padj) cutoff of 0.05 and an absolute log_2_ fold-change > 0.35 to define DEGs. Fig. 7E: We calculated the % common up-, down-regulated genes relative to all DEG in KO-*Dab1*/control and *KO-Cdk5r1-MADM*/control respectively and plotted the mean of these 2 values. Fig. 7F: Significance of the overlap between DEG groups was calculated using newGOM from package GeneOverlap (v1.22.0) using the number of informative genes in this comparison as genome.size. We used different gene groups for further analysis: Common_up defines genes that are common up-regulated DEGs (intersection Fig. 7D left), Common_down defines genes that are common down-regulated DEGs (intersection Fig. 7D right), Dab1_spec defines genes that are DEG in *Dab1*-KO/control but not in *KO-Cdk5r1-MADM*/control (*Dab1* up, *Dab1* down in Fig. 7D) and *Cdk5r1*_spec defines genes that are DEG in *KO-Cdk5r1-MADM*/control but do not overlap (*Cdk5r1* up, *Cdk5r1* down Fig. 7D). GO term enrichment for the respective gene group was performed using enrichGO with parameters: enrichGO(OrgDb = org.Mm.eg.db, readable=T, pool=T, maxGSSize = 900, minGSSize = 100, pvalueCutoff = 0.05, qvalueCutoff = 0.1, separately for different GO ontologies. For Fig. 7G we calculated a score as in Fig. 7E and plotted the top 10 GO terms from cellular components (CC) ontology without prior filtering. For Figure S7 we used the used top 50 GO terms (ranked by adjusted p-value) from *Dab1*_spec and *Cdk5r1*_spec analysis and calculated all pairwise semantic similarity goSim from GOSemSim (v2.12.1) package with parameters: measure="Jiang". We plotted the resulting similarity matrix using the pheatmap package.

### Sample processing for proteomics

Samples were divided into two batches, each containing 3 control samples and 5 *Cdk5r1*-MADM or KO-*Cdk5r1*-MADM samples. Batch 1, processed on day 1, contained green cells from two individual litters and corresponded to control versus KO-*Cdk5r1*-MADM comparison; batch 2, processed on day 2, contains green from 3 individual litters and was used for the control versus *Cdk5r1*-MADM comparison. Each litter contained both control and either *Cdk5r1*-MADM or KO-*Cdk5r1*-MADM. Protein extraction, tryptic digestion, and peptides cleanup were performed using a TMT-labeling compatible variant of the in-Stage Tips method (iST-NHS-12x kit, Preomics). Briefly, immediately after cell-sorting, collected cells were supplemented with 50 µL LYSE-NHS buffer, boiled for 10 min, then processed according to the manufacturer’s protocol with the following minor modifications: sonication was skipped (the number of cells was low enough that DNA would not be an issue, and this would reduce the chance of proteins loss or samples contamination due to having to use a probe sonicator); and digestion was performed overnight. Prior to TMT labeling, small aliquots of each of the 16 samples were taken and mixed to generate a mixed reference sample. Individual samples were then labeled with TMT-10plex (lot # UL291039, ThermoFisher Scientific), splitting the contents of each TMT vial to label one sample of each batch. Individual samples were combined into 2 TMT-labeled samples, each containing the 8 samples from one batch plus a mixed reference sample. Combined samples were then loaded onto the iST-NHS kit’s cartridges in several steps, washed as per the manufacturer’s protocol, eluted, and dried in a speedvac. Since phospho-peptides were of interest, although the amount of material as determined by a Pierce Quantitative Colorimetric Peptide Assay (Thermo Scientific) was low (∼100 µg/sample), the samples were subjected to phospho-peptides enrichment (MagReSyn Ti-IMAC beads, ReSyn Biosciences) according to manufacturer’s protocol but scaling down the amount of beads, then the flow-throughs were fractionated into 8 fractions using the Pierce High pH Reversed-Phase Peptide Fractionation Kit (ThermoFisher Scientific).

### LC-MS/MS analysis

Samples were dried, redissolved in 0.1% TFA and analyzed by LC-MS/MS on an Ultimate HPLC (ThermoFisher Scientific) coupled to a Q-Exactive HF (ThermoFisher Scientific). Each sample was concentrated over an Acclaim PepMap C18 pre-column (5µm particle size, 0.3mm ID x 5 mm length, ThermoFisher Scientific) then bound to a 50cm EasySpray C18 analytical column (2µm particle size, 75μm ID x 500mm length, ThermoFisher Scientific) and eluted over the following 90min gradient: solvent A, water + 0.1% formic acid; solvent B, 80% acetonitrile in water + 0.08% formic acid; constant 300nL/min flow; B percentage: start, 2%; 70min, 31%; 90min, 44%. Mass spectra were acquired in positive mode with a Data Dependent Acquisition method: FWHM 20s, lock mass 445.12003 m/z; MS1: profile mode, 120,000 resolution, AGC target 3e6, 50ms maximum IT, 380 to 1,500m/z; MS2: top 20, centroid mode, 1 microscan, 60,000 resolution, AGC target 1e5, 100ms maximum IT, 0.7m/z isolation window (no offset), 100m/z fixed first mass, NCE 32, excluding charges 1 and 8 or higher, 60s dynamic exclusion.

### Statistical Analysis of Proteomics

Raw files were searched in MaxQuant 1.6.14.0 against the *Mus musculus* reference proteome from UniProtKB. Fixed cysteine modification was set to H11OC6N. Variable modifications were Oxidation (M), Acetyl (Protein N-term), Deamidation (NQ), Gln->pyro-Glu and Phospho (STY). Match between runs and second peptides search were active. All FDRs were set to 1%. MaxQuant results were further processed in R using in-house scripts, starting from evidence (PSM) tables. Briefly, potential contaminants, reverse database hits, or evidences with null intensity values were excluded. Evidence reporter intensities were scaled to integrated feature intensity, normalised using the Levenberg-Marquardt procedure row-wise, then assembled into peptidoforms (post-translationally modified peptides), summing up intensities per sample. Peptidoform reporter intensities were corrected for TMT lot label impurity values, median normalized, log-transformed, subjected to Variance Stabilizing Normalisation, re-normalised using the Levenberg-Marquardt procedure row-wise, then corrected for TMT batch effect using the Internal Reference Scaling method. Ratios to the average of all either control or *Cdk5r1*-MADM samples (two parallel analyses) were calculated, then protein groups were inferred from peptidoforms. Because the focus was on discovering and quantifying as many protein groups as possible from limiting sample amounts, all groups including those with just one peptide were retained. Protein groups were quantified by averaging the intensity profile of matching peptidoforms (excluding phospho-peptides and counterparts), weighted by the inverse of individual Posterior Error Probabilities, then these values were averaged per sample and P values calculated using a moderated t-test and an F-test (limma package). Significance thresholds were calculated using the Benjamini Hochberg procedure for 10, 20 and 30 % FDR. In addition, protein groups with an absolute log2 ratio smaller than 95% of individual to average reference log2 ratios were excluded. Figures 5G-5H: We plotted the Moderated.t-test:.-log10(Pvalue) against the Ratio:.log2.-.Mean for the respective comparison. All genes that were marked “up, FDR = 10%”, “down, FDR = 10%” were labeled in the volcano plot. For Figure 5I: We used all gene names linked to peptide groups for subsequent analyses. We defined significant DEGs as genes marked “up, FDR = 10%”, “down, FDR = 10%” and calculated GO term enrichments using clusterProfiler (v3.14.3) with parameters: Universe = [all genes in “Genes”], OrgDb = org.Mm.eg.db (v3.10.0), ont = ’CC’, pvalueCutoff = 0.1, minGSSize = 50, maxGSSize = 2000, pool=F, readable = T. We plotted selected terms of the top 15 GO terms, ranked by p-value. Figures 5J and 5K: Comparison to RNA-Seq: We defined RNA-Seq DEGs by using statistics calculated in Figure 5B and extracting genes with and adjusted p-value of < 0.1 and a log2 fold-change >0 (RNA Up) or <0 (RNA Down). DEGs based on proteomics were defined as having a “+” in the “Significant:.FDR=10%.-.full-KO” column and “Ratio:.log2.-.Mean.-.full-KO” < 0 (Protein Down) or >0 (Protein Up). Note that we only used genes that were informative on both RNA-Seq and Proteomics for this analysis. Figure 5L: For GO term analysis we used gene sets commonly up-, and down-regulated in both RNA-Seq and proteomics (intersection Figure 5K). GO term enrichment was calculated using enrichGO with parameters: universe = [all genes informative in RNA-Seq and proteomics], OrgDb = org.Mm.eg.db (v3.10.0), ont = ’CC’, pvalueCutoff = 0.9, minGSSize = 10, maxGSSize = 2000, pool=F, readable = T. We calculated a score as described before with a negative prefix for down regulated genes and plotted the top 10 GO terms, ranked by p-value.

## DATA AVAILABILITY

All data have been presented in Figures and Supplemental Figures. Original images will be made available upon request. Raw sequencing/proteomics data, sample lists and code will be accessible as detailed below:

### Transcriptomics raw data

The raw sequencing data used in this publication will be deposited in NCBI’s Gene Expression Omnibus (Edgar et al., 2002) and be accessible through GEO.

### Proteomics raw data

The raw proteomics data used in this publication have been deposited on PRIDE.

### Code for analysis of relative distribution of MADM-Labeled neurons

The Python (version >= 3.6) package cell2layer (version 0.2) for analyzing the relative and absolute distances of each manually marked neuron to its layer boundaries along with documentation is available at http://github.com/hippenmeyerlab/cell2layer as open-source software under the GNU General Public License v3.0.

### Code for correction of non-linear local drift in time-lapse images

The Python (version >= 3.6) package undrift (version 0.2) for correcting non-linear, local tissue drift along with documentation is available at http://github.com/hippenmeyerlab/undrift as open-source software under the GNU General Public License v3.0.

## SUPPLEMENTARY MATERIAL

### Supplementary Figure Legends

**Figure S1.**
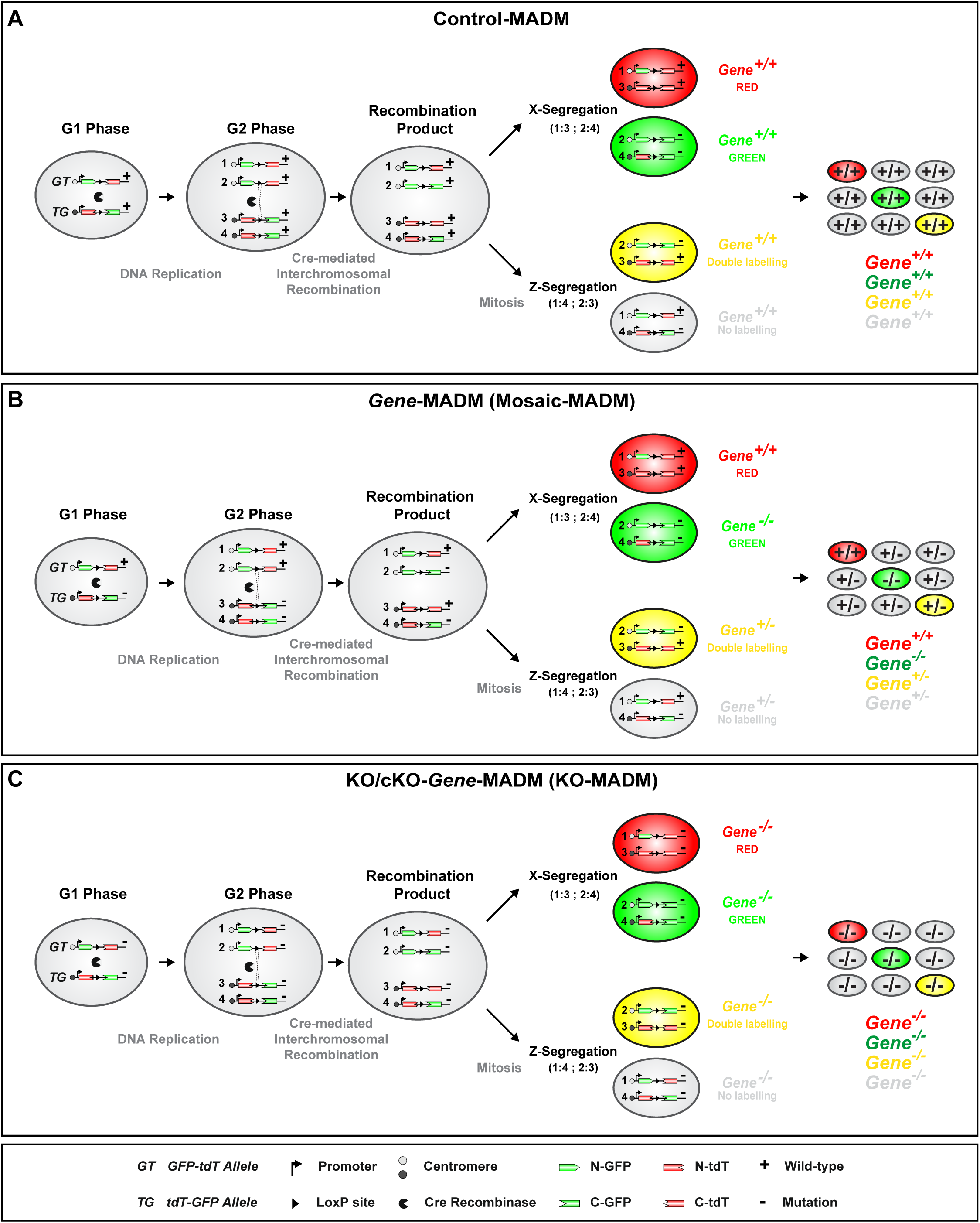
General MADM principle for Control-MADM, Mosaic-MADM and KO/cKO-MADM. **(A)** MADM relies on Cre/loxP-mediated interchromosomal recombination to generate sparse, genetically defined, fluorescently-labeled cells in mice. Thus, it is required that two reciprocal MADM cassettes, each containing the partial N-terminal of the coding sequence for one fluorescent protein (eGFP: enhanced green fluorescent protein) and the C-terminal partial coding sequence for another (tdT: tandem dimer Tomato) separated by an intron containing the loxP-site, are introduced into identical loci on homologous chromosomes. During mitosis, a G2-X event [recombination in G2 of the cell cycle followed by X segregation (top branch)] will result in reconstituted functional green and red fluorescent proteins expressed in each of the two daughter cells, respectively. When G2-Z (bottom branch), [or G1 or G0 (not shown) events] occur, the two fluorescent alleles are passed on together resulting in one double-labeled yellow cell and one unlabeled cell. **(B)** Introduction of a mutation distal to one of the MADM cassettes will allow the generation of genetic mosaics where wild-type (e.g. red) and mutant (e.g. green) daughter cells are each labeled in one color in an otherwise unlabeled heterozygous environment upon G2-X segregation (top branch). In G2-Z (bottom branch), the two fluorescent alleles are passed on together resulting in one double-labeled yellow cell and one unlabeled cell, both heterozygous. **(C)** The introduction of a mutation distal to both of the MADM cassettes allows for the generation of tissue-wide/global KO where all cells are mutant. For more details see (Zong et al 2005; Contreras et al., 2021).

**Figure S2.**
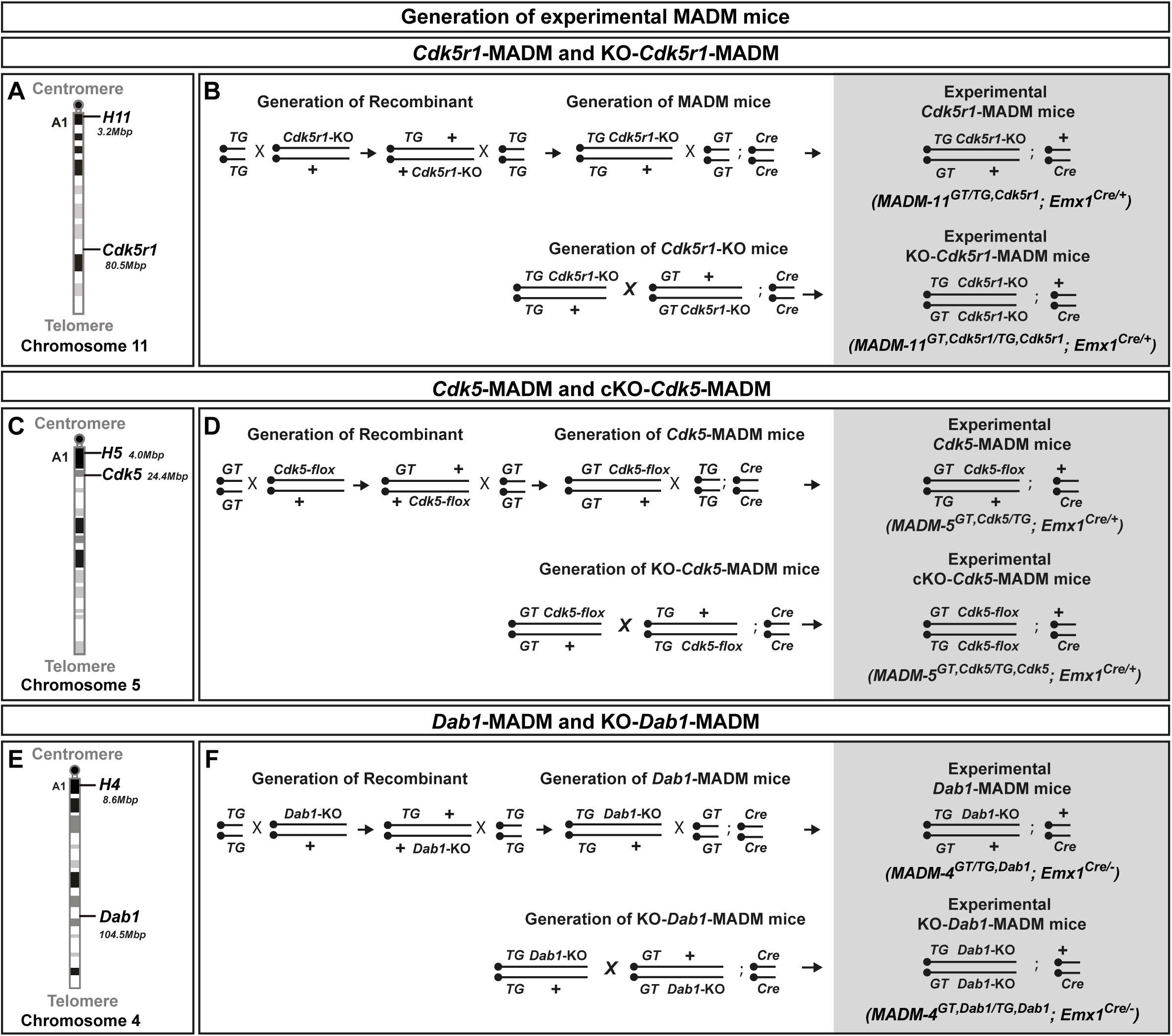
Chromosomal location and MADM breeding schemes for *Cdk5r1*, *Cdk5*, and *Dab1*. **(A)** Location of H11 genomic locus and *Cdk5r1* on mouse chr.11. **(B)** Breeding schemes for the generation of *Cdk5r1-*MADM and KO-*Cdk5r1*-MADM animals. **(C)** Location of H5 genomic locus and *Cdk5* on mouse chr.5. **(D)** Breeding scheme for the generation of *Cdk5-*MADM and KO-*Cdk5*-MADM animals. Note that due to reverse orientation of the MADM cassettes on chr.5, the *Cdk5*-flox allele needs to be recombined to the GT MADM cassette for the generation green labeled *Cdk5^-/-^* mutant cells in mosaic *Cdk5*-MADM (see Contreras et al., 2021 for specific details). **(E)** Location of H4 genomic locus and *Dab1* on mouse chr.4. **(F)** Breeding scheme for the generation of *Dab1-*MADM and KO-*Dab1*-MADM animals.

**Figure S3.**
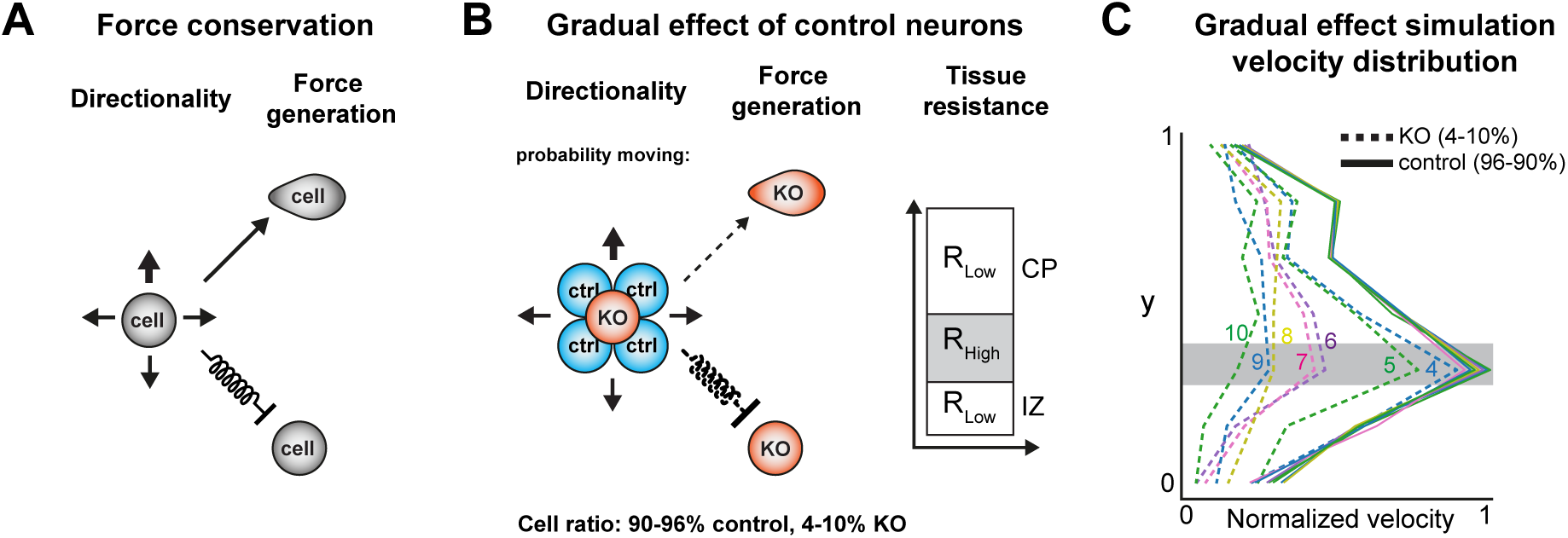
Concepts of modeling approach and gradual effects of mixed model in simulation. **(A)** General concept of model: random walk bias of cells in form of directionality and force generation when cells move and with force conservation included as a spring constant when cells do not move. **(B)** Model setup for mixed neuron migration in Control environment in order to investigate the effect if gradual increase in percentage of Control neurons (ctrl) in mixed population. Simulated Control neurons percentage was varied between 90-96% Control neurons with 4-10% KO. The arrow thickness indicates the probability of the KO neuron to move in any direction. **(C)** Velocity distribution depicting the effect of a gradual increase in percentage of Control cells in mixed population. Numbers indicate the percentage of KO neurons. Note that a cell ratio of less than 4% mutant would yield a control velocity distribution and a cell ratio of more than 10% mutant would yield a KO velocity distribution of the mutant neurons.

**Figure S4.**
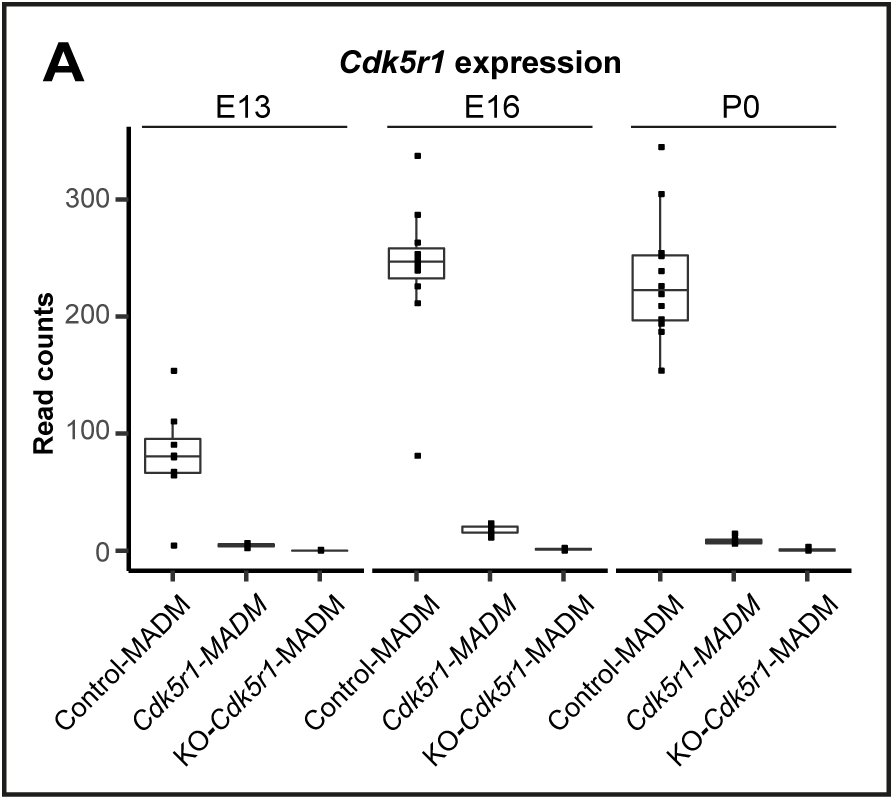
*Cdk5r1* expression levels in MADM-labeled projection neurons at different time points. **(A)** *Cdk5r1* expression (normalized read counts) for each replicate and genotype at E13, E16 and P0.

**Figure S5.**
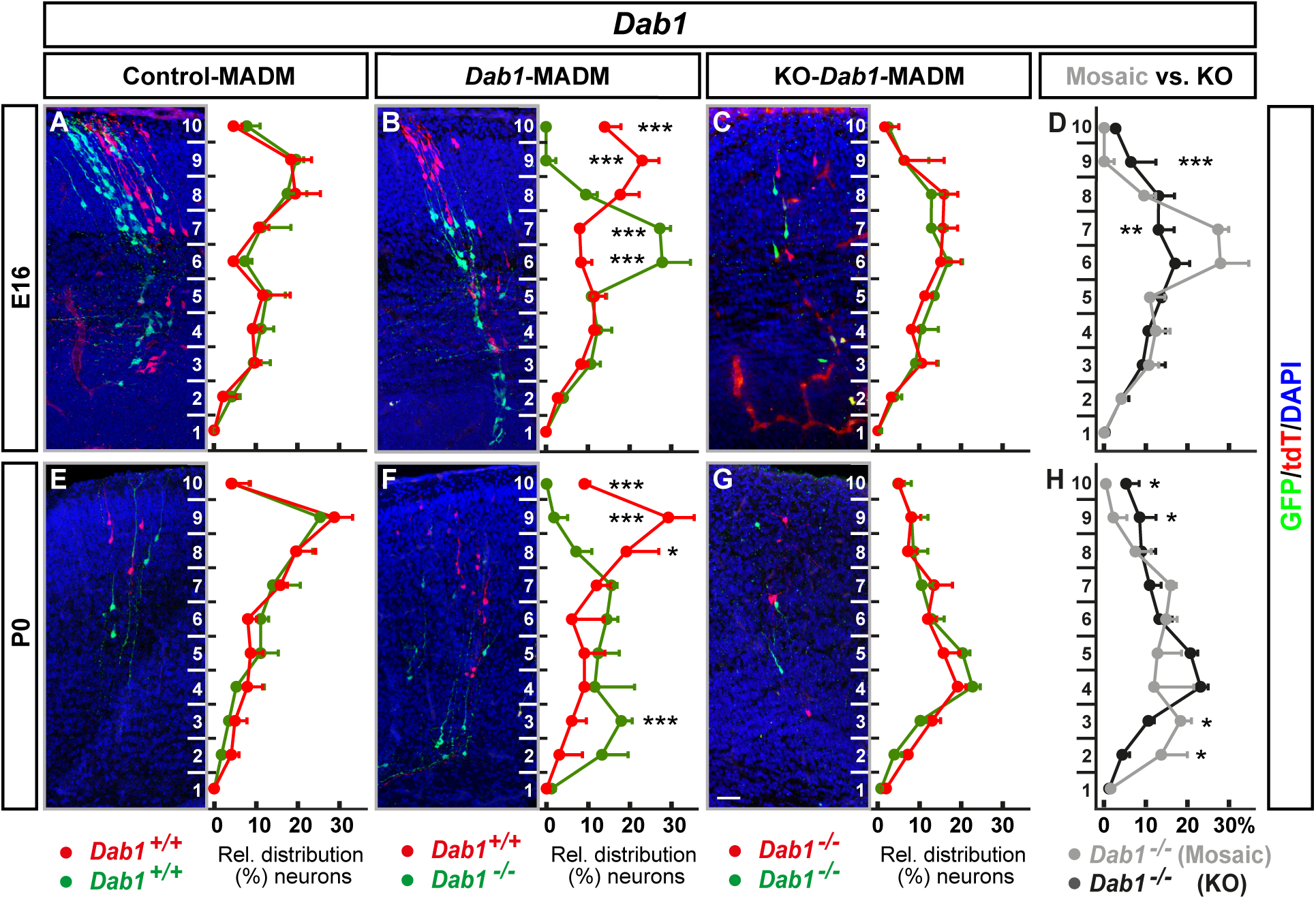
*Dab1* developmental time points E16 & P0 in the somatosensory cortex. **(A-H)** Analysis of green (GFP^+^) and red (tdT^+^) MADM-labeled projection neurons in (A and E) Control-MADM (*MADM-4^GT/TG^;Emx1^Cre/+^*); (B and F) *Dab1*-MADM (*MADM-4^GT/TG,Dab1^;Emx1^Cre/+^*); and (C and G) KO-*Dab1*-MADM (*MADM-4^GT,Dab1/TG,Dab1^;Emx1^Cre/+^*) at E16 (A-D) and P0 (E-H). Relative distribution (%) of MADM-labeled projection neurons is plotted in ten equal bins across the developing cortical wall. (D and H) Direct distribution comparison of *Dab1^-/-^* mutant cells at E16 (D) and P0 (H) in *Dab1*-MADM (grey) versus KO-*Dab1*-MADM (black) distribution. Nuclei were stained using DAPI (blue). n=3 for each genotype (20 hemispheres were analyzed per animal). Data indicate mean ± SD, *p<0.05, **p<0.01, and ***p<0.001. Scale bar: 50µm.

**Figure S6.**
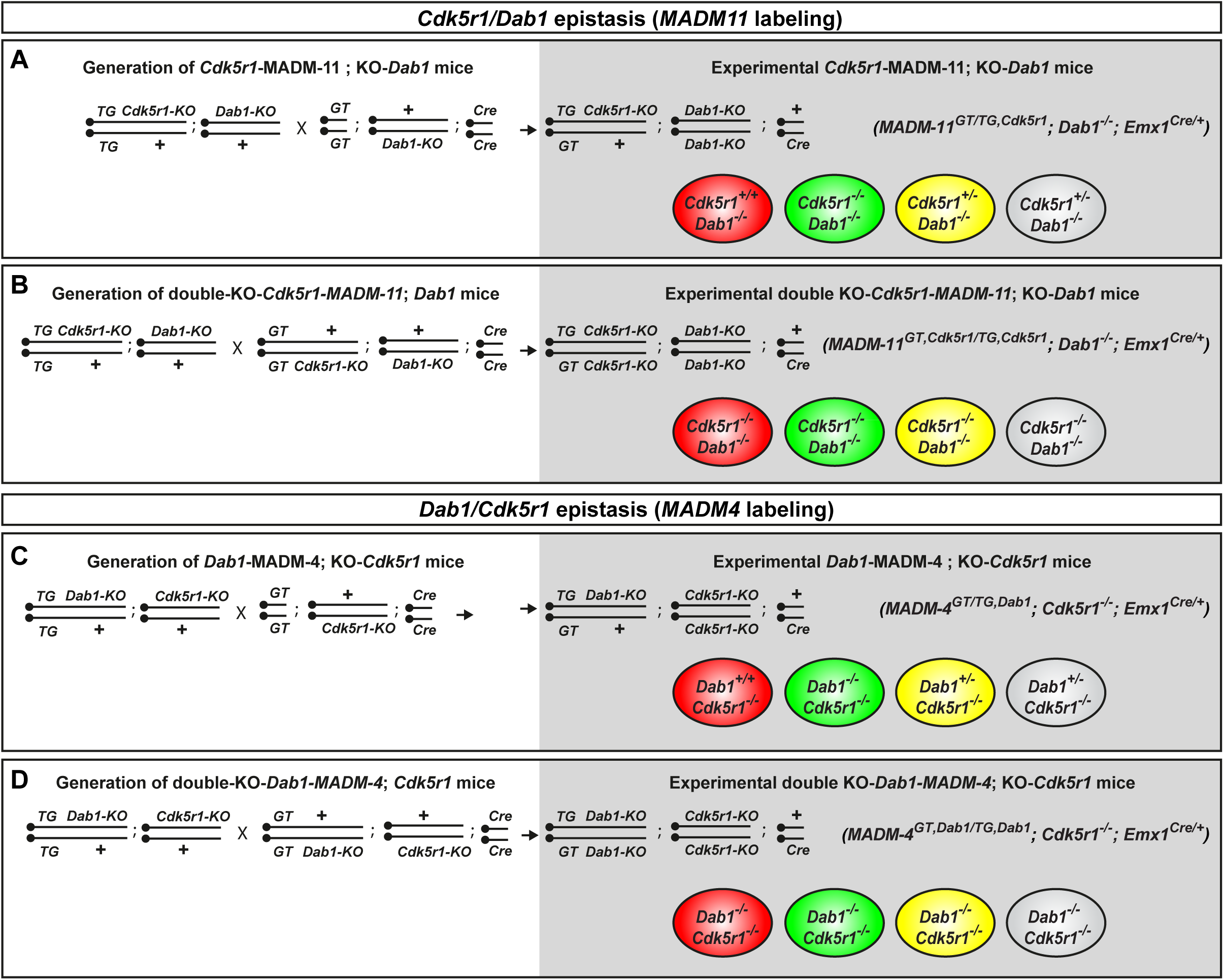
MADM breeding schemes for *Cdk5r1/Dab1* epistasis experiments. **(A-B)** Breeding schemes for *Cdk5r1/Dab1* epistasis experiment using MADM-11 labeling. Generation of experimental (A) mosaic [*Cdk5r1*-MADM-11;KO-*Dab1* (*MADM-11^GT/TG,Cdk5r1^;Dab1^-/-^;Emx1^Cre/+^*)] and (B) double KO [double KO-*Cdk5r1-*MADM-11;KO-*Dab1* (*MADM-11^GT,Cdk5r1/TG,Cdk5r1^;Dab1^-/-^;Emx1-^Cre/+^*)]. **(C-D)** Breeding schemes for *Dab1/Cdk5r1* epistasis experiment using MADM-4 labeling. Generation of experimental (A) mosaic [*Dab1*-MADM-4;KO-*Cdk5r1* (*MADM-4^GT/TG,Dab1^;Cdk5r1^-/-^;Emx1^Cre/+^*)] and (B) double KO [double KO***-****Dab1-*MADM-4;KO-*Cdk5r1* (*MADM-4^GT,Dab1/TG,Dab1^;Cdk5r1^-/-^;Emx1^Cre/+^*)]. Colored cells indicate the respective genotype of the MADM-labeled cells in the experimental animals.

**Figure S7.**
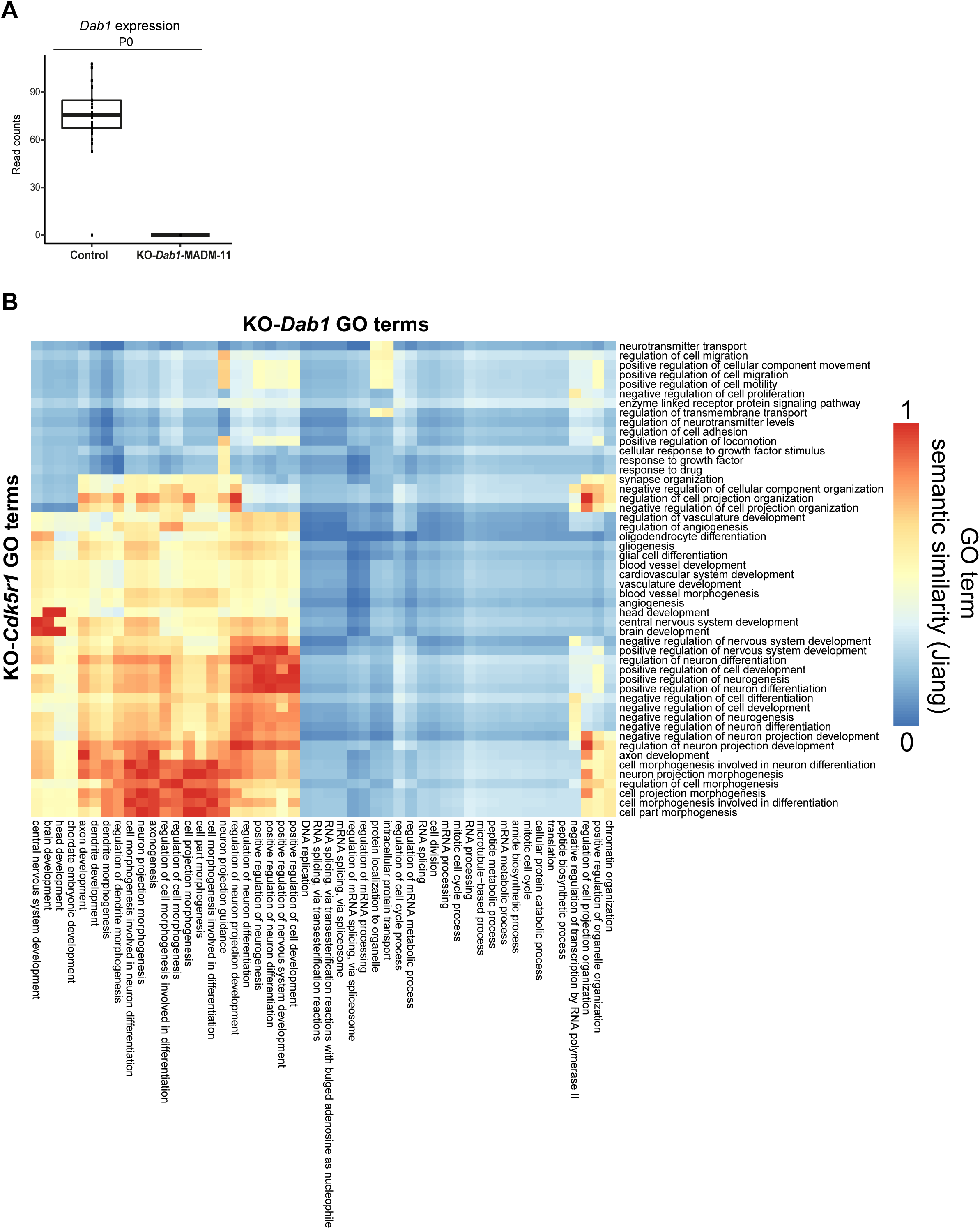
*Dab1* expression and GO term similarity in global *Dab1* and *Cdk5r1* KO. **(A)** *Dab1* expression (normalized read counts) for each replicate analyzed in Figure 7. Control: all *Dab1^+/+^* samples. **(B)** GO term semantic similarity (Jiang) of non-common deregulated genes among GO terms in KO-*Dab1* and KO-*Cdk5r1*, respectively. Similarity value closer to 1 indicates high similarity of the respective GO term.

### Supplementary Video Legends

**Video S1.** Time-lapse imaging of Control-MADM (*MADM-11^GT/TG^;Emx1^Cre/+^*) in somatosensory cortex at E16 during a12h time-period at 15min framerate.

**Video S2.** Time-lapse imaging of *Cdk5r1*-MADM (*MADM-11^GT/TG,Cdk5r1^;Emx1^Cre/+^*) in somatosensory cortex at E16 during a12h time-period at 15min framerate.

**Video S3.** Time-lapse imaging of KO-*Cdk5r1*-MADM (*MADM-11^GT,Cdk5r1/TG,Cdk5r1^; Emx1^Cre/+^*) in somatosensory cortex at E16 during a12h time-period at 15min framerate.

## Notes

### Competing Interest Statement

The authors have declared no competing interest.

